# Multi-omics analysis uncovers differential growth kinetics of Japanese Encephalitis Virus in human and porcine macrophages

**DOI:** 10.1101/2025.06.16.659876

**Authors:** Manas Ranjan Praharaj, Tejaswi Ambati, Sachin Kumar, Baldev Raj Gulati, Subeer S. Majumdar, G. Taru Sharma, Bappaditya Dey, Ravi Kumar Gandham

**Author notes:** Corresponding Authors: G. Taru Sharma, Bappaditya Dey, Ravi Kumar Gandham.

## Abstract

Japanese encephalitis virus (JEV), a mosquito-borne neurotropic virus, poses a significant threat to juvenile humans, while causing mild symptoms in pigs, except piglets. Pigs with high viremia are amplifier hosts, whereas humans with lower viremia are dead-end hosts. The molecular mechanisms behind the varying viremia levels in pigs and humans remain poorly understood. A comparative multi-omics study was conducted on JEV-infected macrophages, the early site of JEV replication in both species. We identified 24 key determinants potentially regulating viral growth. Functional validation of three antiviral factors - LGALS1, ssc-miR-133a-3p, and hsa-miR-3614-5p, and two proviral factors - MIF and VMP1, in macrophages showed that the upregulation of MIF and VMP1, and downregulation of ssc-miR-133a-3p, enhanced JEV replication in pigs. In contrast, the downregulation of MIF and VMP1 and upregulation of hsa-miR-3614-5p reduced JEV replication in human macrophages. These findings provide critical insights into JEV infection and its species-specific dynamics.

## Introduction

Japanese encephalitis virus (JEV), a prominent pathogen belonging to the genus Orthoflavivirus, is prevalent in Southeast Asia and countries such as India, China, and Japan. The malady caused by JEV is encephalitis, which has dire consequences for younglings. JEV is maintained in an enzootic cycle involving herons and egrets as reservoirs, pigs as sentinel and amplifier hosts, and humans and horses as dead-end hosts (Le Flohic et al., 2013). Humans develop low viremia, which is insufficient for mosquito (Culex sp.) transmission, reinforcing their role as dead-end hosts (Le Flohic et al., 2013). The JEV initially replicates in local lymph nodes, keratinocytes, dendritic cells, Langerhans cells, monocytes, and macrophages before spreading to the central nervous system (CNS). Monocytes/macrophages and possibly T-lymphocytes play a crucial role in viral replication and dissemination (Turtle & Solomon, 2018). JEV exploits macrophages for early replication and induces M1 polarization in murine peritoneal macrophages (Wang et al., 2020) and also other studies depicts the role of macrophages during JEV infection (Sooryanarain et al., 2012, Kundu et al., 2013). Given their role in JEV pathogenesis, macrophages were selected in this study to examine viral kinetics in different hosts.

Host factors play a critical role in JEV replication and pathogenesis. A CRISPR-based screen in PK-15 cells identified key regulators of intracellular calcium homeostasis, including CALR, EMC3 (an endoplasmic reticulum membrane complex protein), and proteins involved in the Heparan Sulfate Proteoglycan (HSPG) synthesis pathway (SLC35B2, HS6ST1, B3GAT3, and GLCE), which promote JEV replication in pigs (Zhao et al., 2020). Similarly, a study in human A549 cells identified eleven essential host genes required for viral infection, including PRLHR, ACSL3, and ASCC3 (Liu et al., 2024). Additional studies have shown that host factors, including immune regulators, influence viral replication. Gracia-Nicolas et al. (2019) reported that reduced IFN-β induction renders pig monocyte-derived dendritic cells (MoDCs) more susceptible to JEV than human MoDCs. However, beyond IFN-β induction, differences in host factors likely drive the variation in JEV replication between species.

Despite these findings, it remains unclear whether these host factors account for the differential replication of JEV in amplifying hosts versus dead-end hosts. Addressing this gap is essential for identifying key determinants responsible for variations in viral titers across species. In this study, we used the live attenuated JEV strain (SA14-14-2) due to its replication pattern closely resembling that of the virulent strain (Yun et al., 2016) and its suitability for handling in a BSL-2 laboratory setting. To investigate host-specific differences, we performed proteomics and miRNomics analysis on JEV-infected THP-1-derived macrophages (human) and 3D4/31 cells (pig), aiming to identify host factors and molecular pathways contributing to variations in viral replication.

## Results

### Dynamics of JEV infection in human and porcine macrophages

Human monocyte-derived macrophages infected with JEV at an MOI of 7 (Fig. 1A) exhibited distinct temporal patterns of viral gene and protein expression. NS3 gene expression progressively declined from 24hpi to 36hpi, with no significant change between 36hpi and 48hpi (Fig. 1B). In contrast, NS1 protein expression varied across time points, as observed through immunoblotting (Fig. 1C & 1D). Flow cytometry analysis showed that the percentage of NS1-positive cells was highest at 24hpi and 48hpi, reaching approximately 17–18% (Fig. 1E). Due to uncertainty in deciding the optimal sampling time points for omics studies by these experiments, further studies on cytokine profiling across timepoints were taken up. Secretion of key proinflammatory cytokines (IL-1β, TNF-α, and IL-6), as a measure of host-directed response, was analyzed. TNF-α levels peaked at 36hpi compared to 24hpi and 48hpi (Fig. 1F), while IL-1β and IL-6 levels were significantly higher at 36hpi and 48hpi than at 24hpi (Fig. 1G & 1H). Based on these findings, 36hpi and 48hpi were selected as optimal time points for further experiments in THP-1 cells.

**Figure 1.**
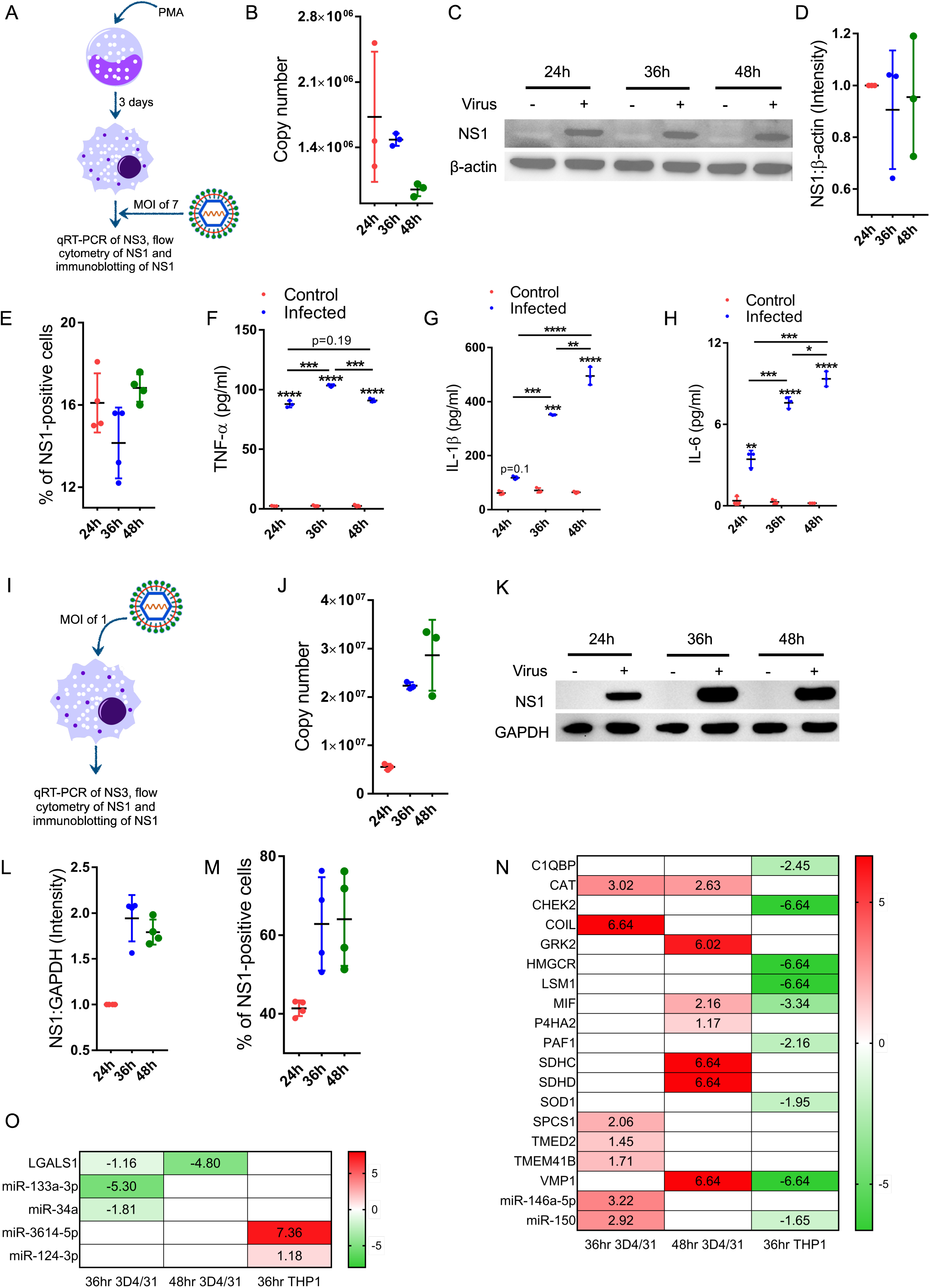
Determination of optimum time points for proteomics and miRNomics. (A) Workflow of THP-1 infection followed by qRT-PCR, immunoblotting, and flow cytometry. (B) qRT-PCR of NS3 gene of JEV between 24hpi, 36hpi and 48hpi in THP-1 and copy number of NS3 was plotted in y-axis. The data represented here were from three independent biological replicates. Error bars represent the mean with SD. (C & D) Immunoblotting of NS1 protein for 24h, 36h, and 48h post-infection in THP-1 and intensity plot of ratio of NS1 to β-actin for 24h, 36h, and 48h post-infection in THP-1. The data represented here were from three independent biological replicates. Error bars represent the mean with SD. (E) Flow cytometry analysis of the percentage of NS1-positive cells for 24h, 36h, and 48h post-infection in THP-1. The data represented here were from four independent biological replicates. Error bars represent the mean with SD. (F-H) Cytokine profiling of TNF-α, IL-1β b, and IL-6 of supernatant collected at 24h, 36h, and 48h post-infection from control and infected THP-1. The data represented here were from three independent biological replicates. Error bars represent the mean with SD. Two-way ANOVA with Bonferroni correction. ∗p < 0.05, ∗∗p < 0.01, ∗∗∗p < 0.001, and ∗∗∗∗p < 0.0001. (I) Workflow of 3D4/31 infection followed by qRT-PCR, immunoblotting, and flow cytometry. (J) qRT-PCR of NS3 gene of JEV between 24hpi, 36hpi and 48hpi in 3D4/31 and copy number of NS3 was plotted in y-axis. The data represented here were from three independent biological replicates. Error bars represent the mean with SD. (K & L) Immunoblotting of NS1 protein for 24h, 36h, and 48h post-infection in 3D4/31 and intensity plot of the ratio of NS1 to GAPDH for 24h, 36h, and 48h post-infection in 3D4/31. The data represented here were from four independent biological replicates. Error bars represent the mean with SD. (M) Flow cytometry analysis of the percentage of NS1-positive cells for 24h, 36h, and 48h post-infection in 3D4/31. The data represented here were from four independent biological replicates. Error bars represent the mean with SD. (N) Heatmap depicting the magnitude of expression of different proviral determinants combining DEPs and DEmiRNAs between 3D4/31 and THP-1. (O) Heatmap depicting the magnitude of expression of different antiviral determinants combining DEPs and DEmiRNAs between 3D4/31 and THP-1.

In porcine 3D4/31 macrophages infected at an MOI of 1, NS3 gene expression was highest at 48hpi, followed by 36hpi and 24hpi (Fig. 1J). However, NS1 protein levels showed only minor differences between 36hpi and 48hpi (Fig. 1K & 1L). Flow cytometry analysis revealed that the percentage of NS1-positive cells was highest at 36hpi and 48hpi (∼65%), followed by 24hpi (∼40%) (Fig. 1M). Notably, porcine macrophages exhibited a higher percentage of NS1-positive cells than human macrophages, even at a lower MOI. Based on these observations, 36hpi and 48hpi were selected for subsequent experiments in 3D4/31 cells.

### Proteomic and miRNome profiling reveal divergent adaptability and infectivity in human and pig macrophages

After identifying distinct patterns of viral kinetics in both hosts, we investigated the global proteome in NS1-positive cells compared to control cells at 36hpi and 48hpi. In 3D4/31 cells at 36hpi, control and infected samples formed distinct clusters, reflecting high intra-group similarity and inter-group differences (Supplementary Fig. 1A). At this time point, 506 differentially expressed proteins (DEPs) were identified, including 381 upregulated and 125 downregulated proteins (Supplementary Fig. 1B). The upregulated DEPs were enriched in pathways related to apoptotic signaling, regulation of viral genome replication, and response to viral infection (Supplementary Fig. 1C), while the downregulated DEPs were associated with cholesterol biosynthesis and cell cycle regulation (Supplementary Fig. 1D). At 48hpi, the control and infected samples exhibited a clustering pattern similar to that observed at 36hpi (Supplementary Fig. 1E). A total of 671 DEPs were identified, with 332 upregulated and 339 downregulated proteins (Supplementary Fig. 1F). Notably, the upregulated DEPs were significantly enriched in immune-related processes compared to 36hpi (Supplementary Fig. 1G). Downregulated DEPs at 48hpi were associated with pathways such as cell cycle regulation, peptide metabolism, and ribosome assembly (Supplementary Fig. 1H). In addition to protein analysis, differentially expressed miRNAs (DEmiRNAs) were examined at 36hpi in 3D4/31 cells. NS1-positive and control cells showed distinct clustering patterns (Supplementary Fig. 1I), with 88 dysregulated DEmiRNAs identified-45 upregulated and 43 downregulated (Supplementary Fig. 1J).

In THP-1 cells at 36hpi, apparent clustering of NS1-positive and control cells highlighted intra-group similarity and inter-group differences (Supplementary Fig. 2A). A total of 685 differentially expressed proteins (DEPs) were identified, including 564 downregulated and 121 upregulated proteins (Supplementary Fig. 2B). Upregulated DEPs were enriched in pathways related to the regulation of cytokine stimulus, negative regulation of viral processes, and response to type I interferon (Supplementary Fig. 2C). Downregulated DEPs were associated with pathways such as RNA processing, mitotic cell cycle regulation, and histone H3-K9 acetylation (Supplementary Fig. 2D). At 48hpi, the clustering pattern remained similar to 36hpi, with distinct grouping of NS1-positive and control samples (Supplementary Fig. 2E). A total of 566 DEPs were observed, including 452 upregulated and 114 downregulated proteins (Supplementary Fig. 2F). Upregulated DEPs were enriched in pathways such as response to virus, negative regulation of macrophage colony-stimulating factor, mitochondrial respiratory chain complex assembly, lipid metabolism, and cytochrome complex assembly (Supplementary Fig. 2G). Downregulated DEPs were associated with processes like protein dephosphorylation and regulation of double strand break repair (Supplementary Fig. 2H). Similarly, PCA clustering of dysregulated miRNAs showed distinct separation between NS1-positive and control cells (Supplementary Fig. 2I). A total of 49 miRNAs were differentially expressed, with 29 downregulated and 20 upregulated (Supplementary Fig. 2J).

Since gene ontology analysis did not yield significant insights, a knowledge-based approach was adopted to identify key DEPs influencing differential adaptability in both hosts, focusing on flaviviruses such as DENV, YFV, WNV, ZIKV, and JEV (Supplementary Fig. 3A). After screening (refer to materials and methods), 24 critical factors were identified as key regulators of viral kinetics (Supplementary file 1), with 19 exhibiting proviral activity (Fig. 1N) and 5 displaying antiviral activity (Fig. 1O). For validation, DEPs (MIF, VMP1, and LGALS1) were selected based on their differential expression between hosts, while DEmiRNAs (ssc-miR-133a-3p and hsa-miR-3614-5p) were prioritized due to their previously uncharacterized roles in JEV infection. Modulating the expression of these factors in the opposite direction of their observed patterns during infection further highlighted their regulatory impact. The qRT-PCR, immunoblotting, and plaque assay were performed to check the JEV replication, translation and release, respectively.

### MIF maintains the viral niche and acts as a proviral determinant

MIF (Macrophage Migration Inhibitory Factor) has been reported to play a protective role in JEV infection (Suzuki et al., 2000) and a significant role in maintaining DENV viremia (Assunção-Miranda et al.,2010). MIF, upregulated in 3D4/31 cells but downregulated in THP-1 cells (Fig. 1N), was validated for its proviral role in 3D4/31 cells using a drug inhibition assay with 20 µM ISO-1, following a specified treatment regime (Fig. 2A). This treatment reduced MIF expression at 36 h post-treatment (Fig. 2B). After confirming drug activity, cells were infected with JEV, and samples were collected at 36hpi and 48hpi. qRT-PCR analysis of NS3 showed a significant decrease in NS3 expression at 48hpi in ISO-1-treated cells compared to DMSO-treated controls (Fig. 2C). Additionally, NS1 expression was significantly lower at 36hpi in ISO-1-treated cells than in vehicle control (Fig. 2D & 2E), reinforcing MIF’s proviral role. Immunoblotting for NS3 after MIF depletion further confirmed reduced NS3 protein levels at 36hpi and 48hpi (Fig. 2F & 2G). Finally, a plaque assay demonstrated reduced viral titers in ISO-1-treated cells compared to controls (Fig. 2H). To confirm MIF’s proviral role, ΔMIF 3D4/31 cells were generated and validated by immunoblotting (Fig. 2I). Following JEV infection, qRT-PCR showed a significant reduction in NS3 copy number at 48hpi (Fig. 2J), which was further corroborated by immunoblotting that revealed decreased NS3 expression in MIF-deleted cells (Fig. 2K & 2L). These findings confirm that MIF enhances JEV replication, supporting its role as a proviral factor in 3D4/31 cells.

**Figure 2.**
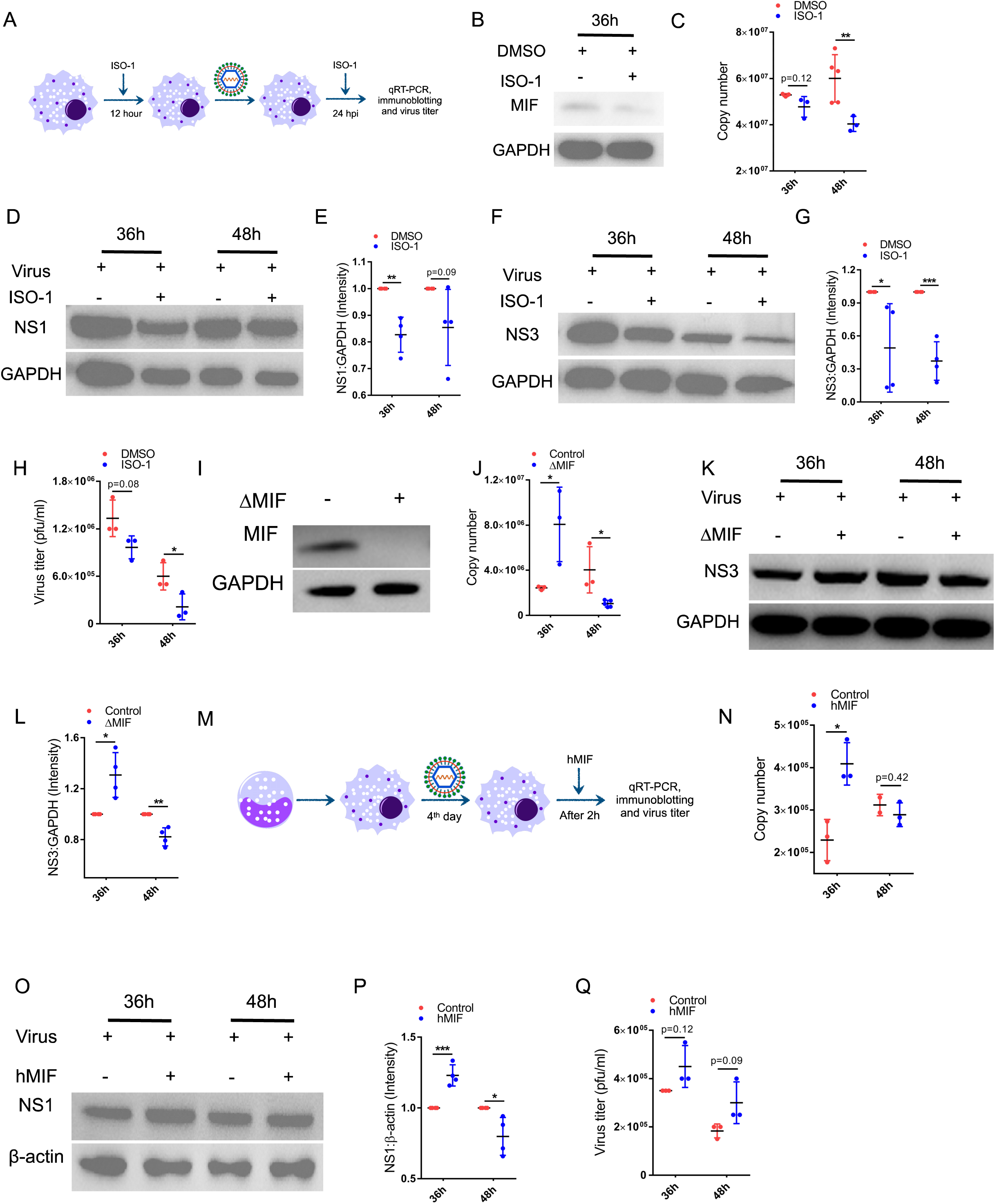
Role of MIF in alteration of virus titer between 3D4/31 and THP-1. (A) Flow chart for the ISO-1 treatment regime carried out to inhibit MIF, followed by qRT-PCR and immunoblotting. (B) Immunoblotting of MIF protein at 36h in 3D4/31 with or without ISO-1 treatment. The data is representative of three independent experiments. (C) qRT-PCR of the NS3 gene of JEV at 36hpi and 48hpi in 3D4/31 after different treatments, and the copy number of NS3 was plotted in the y-axis, representing the differences between DMSO and ISO-1 treated cells. The data represented here were from three independent biological replicates, with two biological replicates repeated one more time for DMSO treatment at 48hpi. Error bars represent the mean with SD. Unpaired t-test. ∗p < 0.05, ∗∗p < 0.01, ∗∗∗p < 0.001, and ∗∗∗∗p < 0.0001. (D & E) Immunoblotting of NS1 protein at 36hpi and 48hpi in 3D4/31 and intensity plot of the ratio of NS1 to GAPDH for 36hpi and 48hpi in 3D4/31 after different treatment regimes. Data represented here were from four independent biological replicates. Error bars represent the mean with SD. Unpaired t-test. ∗p < 0.05, ∗∗p < 0.01, ∗∗∗p < 0.001, and ∗∗∗∗p < 0.0001. (F & G) Immunoblotting of NS3 protein at 36hpi and 48hpi in 3D4/31 and intensity plot of the ratio of NS3 to GAPDH for 36hpi and 48hpi in 3D4/31 after different treatment regimes. The data represented here were from four independent biological replicates. Error bars represent the mean with SD. Unpaired t-test. ∗p < 0.05, ∗∗p < 0.01, ∗∗∗p < 0.001, and ∗∗∗∗p < 0.0001. (H) Plaque assay to determine the virus titer compared between ISO-1 and DMSO treatment at 36hpi and 48hpi. The data represented here were from three independent biological replicates. Error bars represent the mean with SD. Unpaired t-test. ∗p < 0.05, ∗∗p < 0.01, ∗∗∗p < 0.001, and ∗∗∗∗p < 0.0001. (I) Immunoblotting of MIF protein in 3D4/31 Cas9 and 3D4/31 ΔMIF. The data is representative of three independent experiments. (J) qRT-PCR of NS3 gene of JEV at 36hpi and 48hpi in 3D4/31 Cas9 and 3D4/31 ΔMIF and copy number of NS3 was plotted in y-axis representing the differences between 3D4/31 Cas9 and 3D4/31 ΔMIF. The data represented here were from three independent replicates, except for five independent replicates from 48hpi 3D4/31 ΔMIF. Error bars represent the mean with SD. Unpaired t-test. ∗p < 0.05, ∗∗p < 0.01, ∗∗∗p < 0.001, and ∗∗∗∗p < 0.0001. (K & L) Immunoblotting of NS3 protein at 36hpi and 48hpi in 3D4/31 Cas9 and 3D4/31 ΔMIF and intensity plot of the ratio of NS3 to GAPDH for 36hpi and 48hpi in 3D4/31 Cas9 and 3D4/31 ΔMIF. The data represented here were from two independent biological replicates. Error bars represent the mean with SD. Unpaired t-test. ∗p < 0.05, ∗∗p < 0.01, ∗∗∗p < 0.001, and ∗∗∗∗p < 0.0001. (M) A flow chart for the human recombinant MIF treatment regime was carried out to increase the MIF level in THP-1. (N) qRT-PCR of NS3 gene of JEV at 36hpi and 48hpi in THP-1 after hMIF treatment, and the copy number of NS3 was plotted in the y-axis, representing the differences between control and hMIF-treated cells. The data represented here were from three independent biological replicates, except for 48hpi control cells, which had two independent biological replicates. Error bars represent the mean with SD. Unpaired t-test. ∗p < 0.05, ∗∗p < 0.01, ∗∗∗p < 0.001, and ∗∗∗∗p < 0.0001. (O & P) Immunoblotting of NS1 protein at 36hpi and 48hpi in THP-1 and intensity plot showing the ratio of NS1 to β-actin for 36hpi and 48hpi in THP-1 after hMIF treatment. The data represented here were from two independent biological replicates. Error bars represent the mean with SD. Unpaired t-test. ∗p < 0.05, ∗∗p < 0.01, ∗∗∗p < 0.001, and ∗∗∗∗p < 0.0001. (Q) Plaque assay to determine the virus titer compared between control and hMIF treatment at 36hpi and 48hpi. Data represented here were from three independent biological replicates. Error bars represent the mean with SD. Unpaired t-test. ∗p < 0.05, ∗∗p < 0.01, ∗∗∗p < 0.001, and ∗∗∗∗p < 0.0001.

Similarly, in THP-1 cells, recombinant MIF supplementation (Fig. 2M) increased NS3 gene expression at 36hpi (Fig. 2N) and elevated NS1 protein levels (Fig. 2O & 2P). Although plaque assays suggested enhanced virus release, the increase was insignificant (Fig. 2Q). These findings establish MIF as a key regulator of JEV infection, influencing viral replication kinetics in a host-dependent manner.

### VMP1 facilitates JEV replication in human and porcine macrophages

VMP1, known for its roles in autophagy and lipid mobilization (Morishita et al., 2019, Ropolo et al., 2007, Zhao et al., 2017), has been implicated in facilitating DENV replication (Yousefi et al., 2022, Hoffmann et al., 2021). VMP1 was upregulated at 48hpi in 3D4/31 cells but downregulated at 36hpi in THP-1 cells (Fig. 1N), suggesting a potential role in facilitating JEV replication. To evaluate VMP1’s role in JEV infection, its expression was silenced in 3D4/31 cells using siRNAs (Fig. 3A). qRT-PCR confirmed the effective knockdown of VMP1 (Fig. 3B). Following JEV infection, decrease in NS3 gene expression indicated decrease in viral RNA at 36hpi in VMP1-silenced cells (Fig. 3C). Further, a significant decrease in NS1 protein levels at 36 hpi on immunoblotting (Fig. 3D & 3E) along with a lower percentage of NS1-positive cells at 48hpi (Fig. 3F) confirmed the role of VMP1 in facilitating JEV replication. In addition, an observed, though not significant, reduction in virus release was seen at 36hpi after VMP1 knockdown (Fig. 3G).

**Figure 3.**
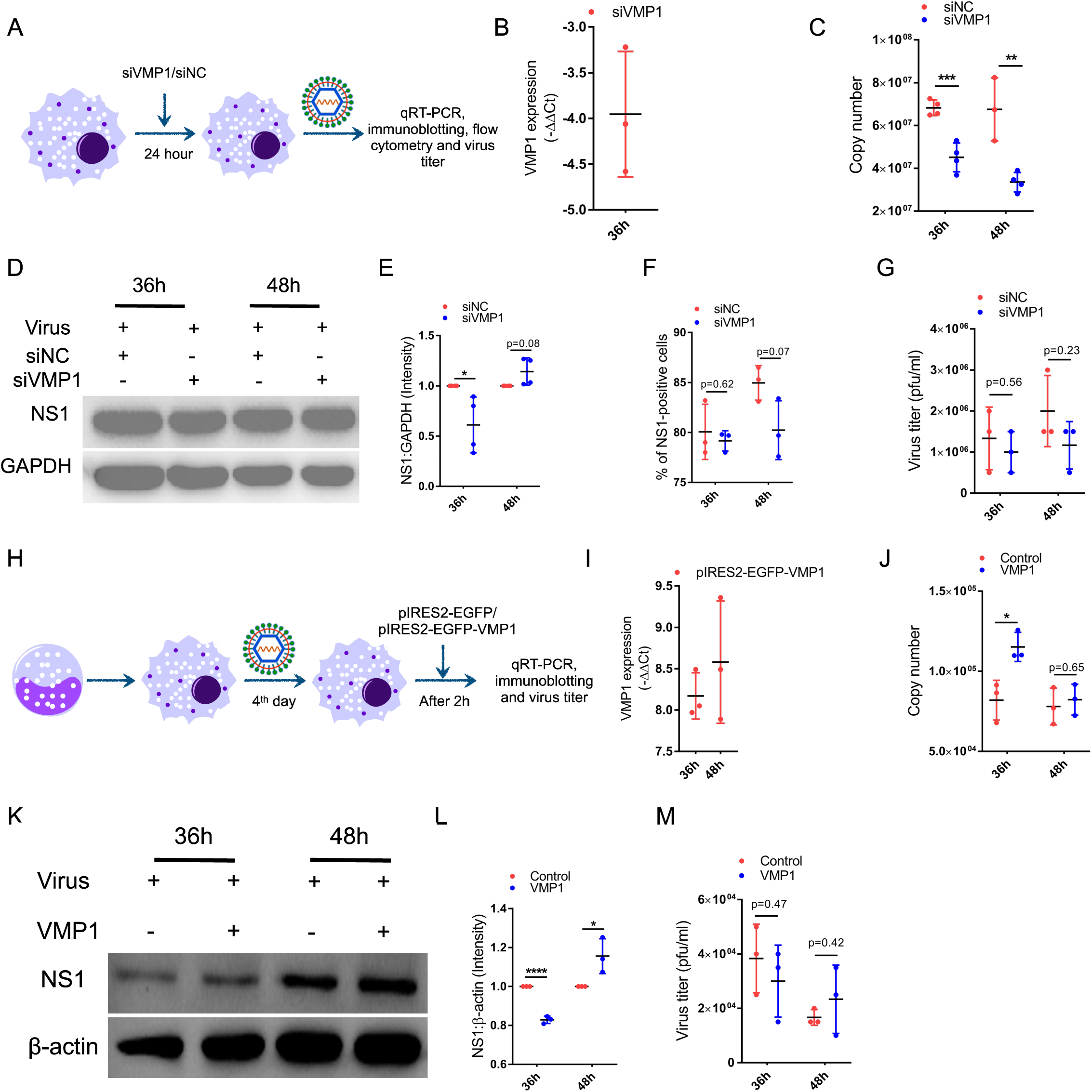
VMP1 as an aid in maintaining the viral niche in pig and human macrophages. (A) Flow chart for the siNC/SiVMP1 treatment regime carried out to inhibit VMP1, followed by qRT-PCR and immunoblotting. (B) qRT-PCR of VMP1 gene at 36h in 3D4/31 with or without siVMP1 treatment. The data represented here were from three independent biological replicates. Error bars represent the mean with SD. Unpaired t-test. ∗p < 0.05, ∗∗p < 0.01, ∗∗∗p < 0.001, and ∗∗∗∗p < 0.0001. (C) qRT-PCR of NS3 gene of JEV at 36hpi and 48hpi in 3D4/31 after siNC or siVMP1 treatment, and the copy number of NS3 was plotted on the y-axis, representing the differences between siNC and siVMP1-treated cells. The data represented here were from three independent biological replicates, with one biological replicate repeated one more time except for siNC treatment at 48hpi. Error bars represent the mean with SD. Unpaired t-test. ∗p < 0.05, ∗∗p < 0.01, ∗∗∗p < 0.001, and ∗∗∗∗p < 0.0001. (D & E) Immunoblotting of NS1 protein at 36hpi and 48hpi in 3D4/31 and intensity plot showing the ratio of NS1 to GAPDH for 36hpi and 48hpi in 3D4/31 after siNC or siVMP1 treatment. Data represented here were from four independent biological replicates. Error bars represent the mean with SD. Unpaired t-test. ∗p < 0.05, ∗∗p < 0.01, ∗∗∗p < 0.001, and ∗∗∗∗p < 0.0001. (F) Flow cytometry analysis of the percentage of NS1-positive cells at 36hpi and 48hpi in 3D4/31 after siNC or siVMP1 treatment. The data represented here were from three independent biological replicates. Error bars represent the mean with SD. Unpaired t-test. ∗p < 0.05, ∗∗p < 0.01, ∗∗∗p < 0.001, and ∗∗∗∗p < 0.0001. (G) Plaque assay to determine the virus titer compared between siNC or siVMP1 treatment at 36hpi and 48hpi. The data represented here were from three independent biological replicates. Error bars represent the mean with SD. Unpaired t-test. ∗p < 0.05, ∗∗p < 0.01, ∗∗∗p < 0.001, and ∗∗∗∗p < 0.0001. (H) Flow chart for the pIRES2-EGFP or pIRES2-EGFP-VMP1 plasmid transfection regime carried out to overexpress VMP1, followed by qRT-PCR and immunoblotting. (I) qRT-PCR of VMP1 gene at 36hpi and 48hpi in THP-1 with pIRES2-EGFP or pIRES2-EGFP-VMP1 plasmid. The data represented here were from three independent biological replicates. (J) qRT-PCR of NS3 gene of JEV at 36hpi and 48hpi in THP-1 after pIRES2-EGFP or pIRES2-EGFP-VMP1 plasmid transfection and the copy number of NS3 was plotted in the y-axis, representing the differences between pIRES2-EGFP or pIRES2-EGFP-VMP1 plasmid-transfected cells. The data represented here were from three independent biological replicates. Error bars represent the mean with SD. Unpaired t-test. ∗p < 0.05, ∗∗p < 0.01, ∗∗∗p < 0.001, and ∗∗∗∗p < 0.0001. (K & L) Immunoblotting of NS1 protein at 36hpi and 48hpi in THP-1 and intensity plot showing the ratio of NS1 to GAPDH for 36hpi and 48hpi in THP-1 after pIRES2-EGFP or pIRES2-EGFP-VMP1 plasmid transfection. The data represented here were from three independent biological replicates. Error bars represent the mean with SD. Unpaired t-test. ∗p < 0.05, ∗∗p < 0.01, ∗∗∗p < 0.001, and ∗∗∗∗p < 0.0001. (M) Plaque assay to determine the virus titer compared between pIRES2-EGFP and pIRES2-EGFP-VMP1 plasmid transfected cells at 36hpi and 48hpi. The data represented here were from three independent biological replicates. Error bars represent the mean with SD. Unpaired t-test. ∗p < 0.05, ∗∗p < 0.01, ∗∗∗p < 0.001, and ∗∗∗∗p < 0.0001.

To investigate VMP1’s role in JEV infection in THP-1 cells, pIRES2-EGFP or pIRES2-EGFP-VMP1 plasmid was transfected 2 hours after viral overlay (Fig. 3H). qRT-PCR confirmed a significant increase in VMP1 expression following plasmid transfection (Fig. 3I). Transient VMP1 expression led to a notable rise in NS3 gene expression at 36 hpi (Fig. 3J). Immunoblotting further confirmed increased NS1 protein levels at 48 hpi in pIRES2-EGFP-VMP1-transfected cells (Fig. 3K & 3L). Moreover, the virus release was found to increase at 48hpi upon VMP1 expression, though not significant (Fig. 3M). These findings suggest that VMP1 enhances JEV replication in THP-1 cells.

### LGALS1 exhibits limited influence on JEV replication in macrophages

Gal-1 attenuates virus adsorption and internalisation in case of DENV infection, further limiting virus egress (Toledo et al., 2014). In this study, LGALS1 expression was downregulated in pig 3D4/31 cells, in contrast to its upregulation in THP-1-derived macrophages (not significant). LGALS1 expression was found to be downregulated in 3D4/31 cells. To assess its role in JEV replication, we generated a stable cell line overexpressing LGALS1 fused with mCherry, which was confirmed via immunoblotting (Fig. 4A). Following this confirmation, we infected the cells and observed that NS3 expression was reduced in LGALS1-overexpressing cells at 48 h post-infection (hpi). However, the reduction was not statistically significant (Fig. 4B). Interestingly, immunoblotting for NS1 revealed no significant difference in expression levels between LGALS1-overexpressing and control cells (Fig. 4C). Moreover, the virus release was seen to decrease at 48hpi in LGALS1-overexpressing cells. Still, the decrease was not significant (Fig. 4D). Collectively, these findings suggest that LGALS1 overexpression does not substantially influence viral replication or kinetics in 3D4/31 cells, indicating it likely plays no significant role in maintaining the JEV replication niche in this cell type.

**Figure 4.**
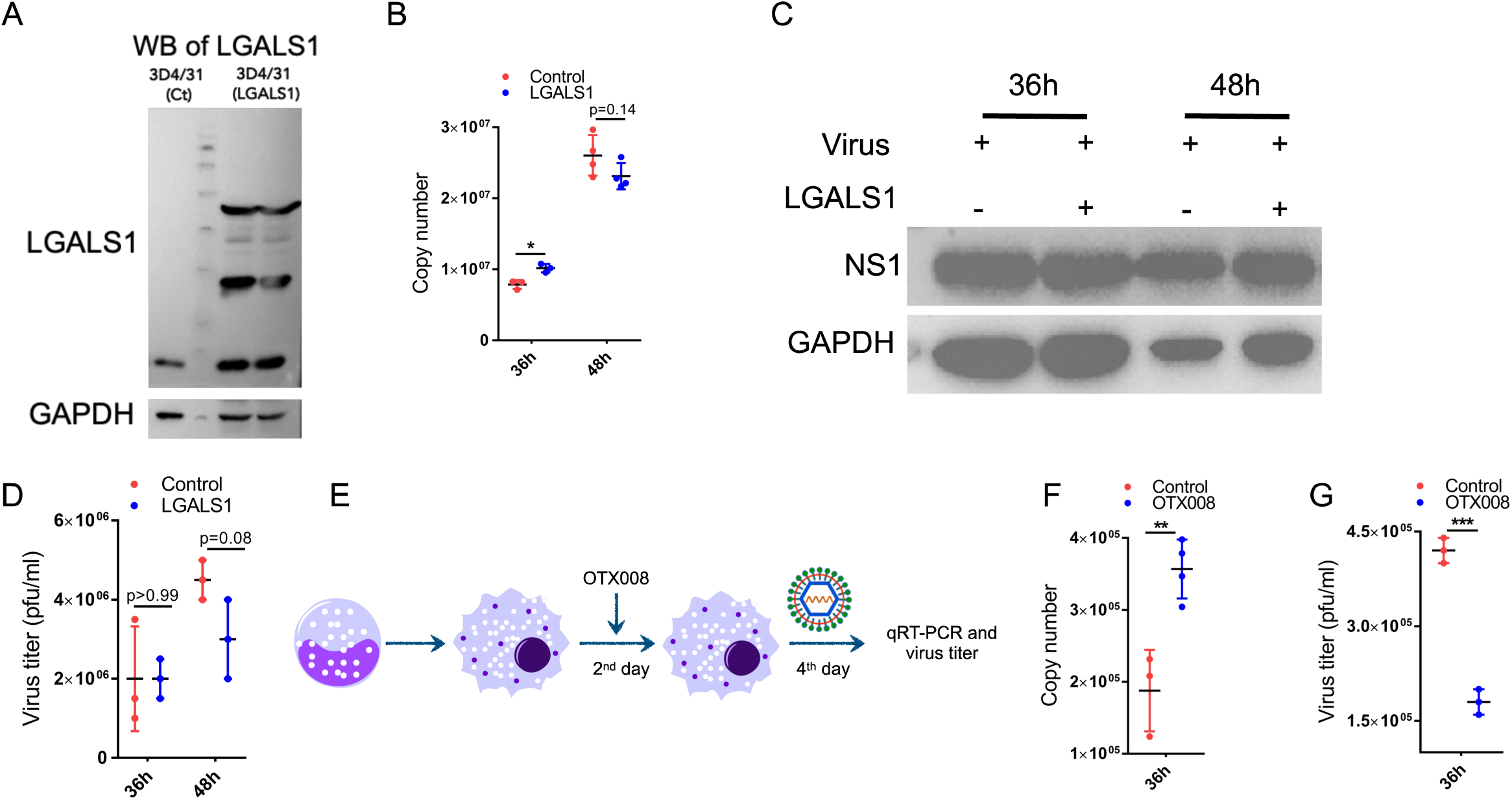
LGALS1 has a minuscule role in maintaining the viral niche. (A) Immunoblotting of LGALS1 protein at 36h in 3D4/31 mCherry and 3D4/31 LGALS1. The data is representative of three independent experiments. (B) qRT-PCR of NS3 gene of JEV at 36hpi and 48hpi in 3D4/31 mCherry and 3D4/31 LGALS1 and copy number of NS3 was plotted in y-axis representing the differences between 3D4/31 mCherry and 3D4/31 LGALS1 cells. Data represented at 36h were from three independent biological replicates, and at 48h were from two independent biological replicates. Error bars represent the mean with SD. Unpaired t-test. ∗p < 0.05, ∗∗p < 0.01, ∗∗∗p < 0.001, and ∗∗∗∗p < 0.0001. (C) Immunoblotting of NS1 protein at 36hpi and 48hpi in 3D4/31 mCherry and 3D4/31 LGALS1 cells. The data is representative of three independent experiments. (D) Plaque assay to determine the virus titer in 3D4/31 mCherry and 3D4/31 LGALS1 cells at 36hpi and 48hpi. The data represented here were from three independent biological replicates. Error bars represent the mean with SD. Unpaired t-test. ∗p < 0.05, ∗∗p < 0.01, ∗∗∗p < 0.001, and ∗∗∗∗p < 0.0001. (E) Flow chart for the OTX008 treatment regime carried out to inhibit LGALS1. (F) qRT-PCR of NS3 gene of JEV at 36hpi in THP-1 and OTX008-treated THP-1, and the copy number of NS3 was plotted in the y-axis, representing the differences between THP-1 and OTX008-treated THP-1 cells. Data represented here were from three independent biological replicates, with one biological replicate being repeated one more time in OTX008-treated cells. Error bars represent the mean with SD. Unpaired t-test. ∗p < 0.05, ∗∗p < 0.01, ∗∗∗p < 0.001, and ∗∗∗∗p < 0.0001. (G) Plaque assay to determine the virus titer compared between THP-1 and OTX008-treated THP-1 at 36hpi. The data represented here were from three independent biological replicates. Error bars represent the mean with SD. Unpaired t-test. ∗p < 0.05, ∗∗p < 0.01, ∗∗∗p < 0.001, and ∗∗∗∗p < 0.0001.

To further explore LGALS1 activity in THP-1 cells during JEV infection, we treated the cells with OTX008, a known LGALS1 inhibitor, followed by infection (Fig. 4E). Surprisingly, NS3 expression was increased in OTX008-treated cells at 36hpi compared to untreated controls (Fig. 4F). However, plaque assays revealed a significant reduction in viral titers in OTX008-treated cells after infection (Fig. 4G). These results suggest that it does not play a critical role in JEV replication dynamics in THP-1 cells.

### ssc-miR-133a-3p binds to 3’UTR of JEV and inhibits virus replication in pig macrophages

The 3′UTR of the flavivirus genome plays a crucial role in viral replication (Jiang et al., 2009). Overexpression of miR-133a-3p in DENV-infected Vero cells has been shown to inhibit viral replication by targeting the 3′UTR (Castillo et al., 2016). In this study, binding between ssc-miR-133a-3p and the JEV 3′UTR was predicted using Bibiserv2, with a calculated binding energy of −25.7 kcal/mol (Supplementary Fig. 4A). A dual-luciferase assay confirmed this interaction, as ssc-miR-133a-3p significantly reduced luciferase activity in pmirGLO-3′UTR-transfected HEK293 cells but had no effect in pmirGLO-only transfected cells (Fig. 5A, Supplementary Fig. 4B).

**Figure 5.**
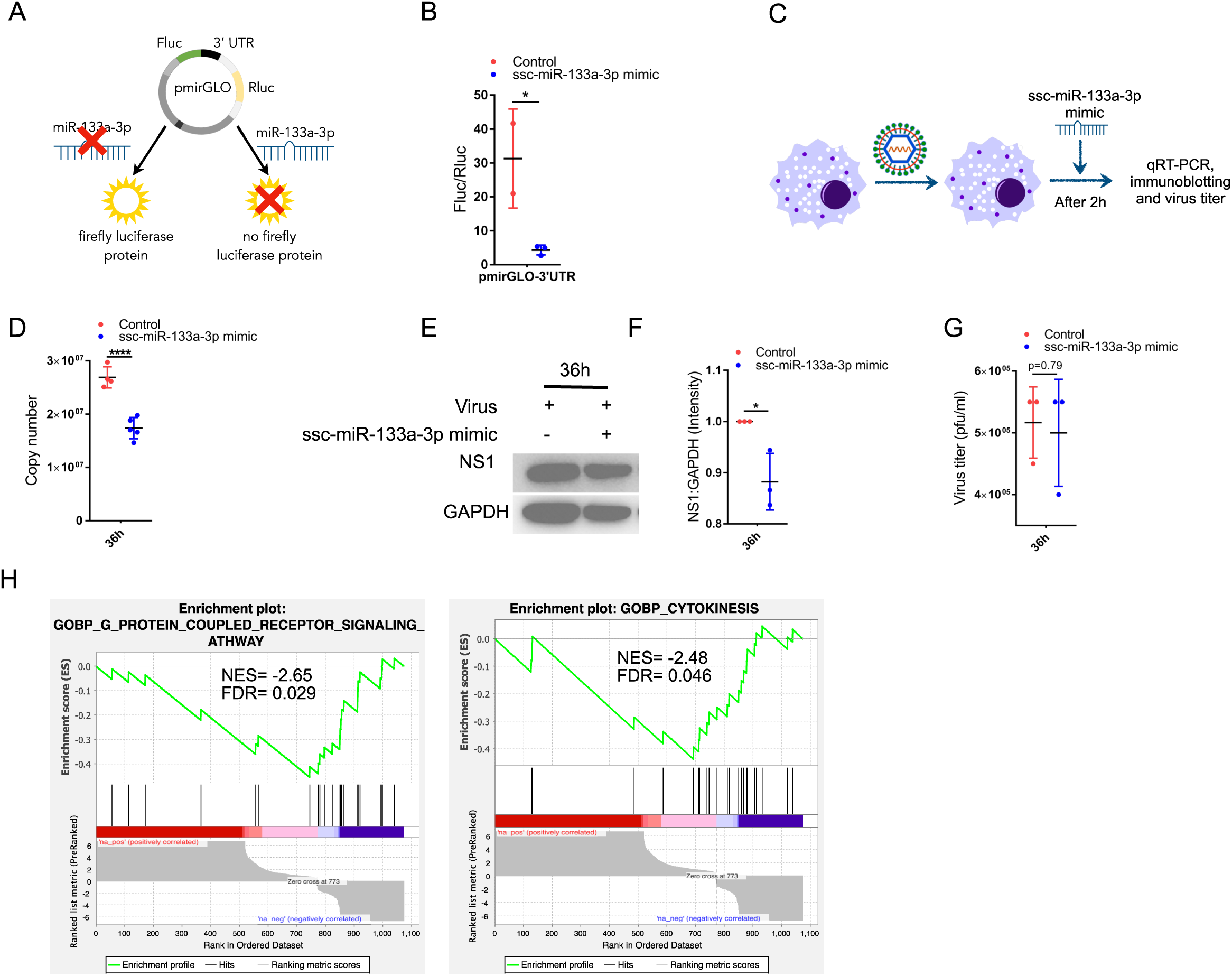
ssc-miR-133a-3p is an antiviral. (A) Schematic representation of dual luciferase assay. (B) The plot represents the ratio of Fluc to Rluc, showing the differences between control and mimic transfected HEK293 transfected with pmirGLO-3’UTR. The data represented here were from two independent biological replicates for control cells and three independent biological replicates for mimic-transfected cells. Error bars represent the mean with SD. Unpaired t-test. ∗p < 0.05, ∗∗p < 0.01, ∗∗∗p < 0.001, and ∗∗∗∗p < 0.0001. (C) Workflow showing the protocol used for mimic transfection in 3D4/31 cells. (D) qRT-PCR of NS3 gene of JEV at 36hpi in 3D4/31 and mimic transfected 3D4/31 cells and copy number of NS3 was plotted in y-axis representing the differences between 3D4/31 and mimic transfected 3D4/31 cells. The data represented here were from three independent biological replicates, with one biological replicate being repeated in control cells and two biological replicates being repeated one more time in mimic-transfected cells. Error bars represent the mean with SD. Unpaired t-test. ∗p < 0.05, ∗∗p < 0.01, ∗∗∗p < 0.001, and ∗∗∗∗p < 0.0001. (E & F) Immunoblotting of NS1 protein at 36hpi in 3D4/31 and mimic-transfected 3D4/31 cells and intensity plot of the ratio of NS1 to GAPDH of 36hpi in 3D4/31 and mimic-transfected 3D4/31 cells. The data represented here were from three independent biological replicates. Error bars represent the mean with SD. Unpaired t-test. ∗p < 0.05, ∗∗p < 0.01, ∗∗∗p < 0.001, and ∗∗∗∗p < 0.0001. (G) Plaque assay to determine the virus titer compared between 3D4/31 and mimic-transfected 3D4/31 cells at 36hpi. The data represented here were from three independent biological replicates. Error bars represent the mean with SD. Unpaired t-test. ∗p < 0.05, ∗∗p < 0.01, ∗∗∗p < 0.001, and ∗∗∗∗p < 0.0001. (H) GSEA enrichment plot showing the enrichment score of biological processes using the pre-ranked file between mimic-transfected and infected cells vs. infected cells.

To assess its functional impact, ssc-miR-133a-3p mimic was transfected into JEV-infected 3D4/31 cells (Fig. 5C) and confirmed for increased miRNA expression by qRT-PCR (Supplementary Fig. 4C). On JEV infection and transfection, at 36hpi NS3 gene expression was reduced (Fig. 5D) accompanied by lower NS1 protein levels (Fig. 5E & 5F), indicating suppression of viral replication and translation. Plaque assays showed a reduction in virus titers at this time, which was insignificant (Fig. 5G). These findings suggest that ssc-miR-133a-3p exerts an antiviral effect by targeting the JEV 3′UTR to suppress replication and translation.

In addition to directly targeting the 3′UTR, ssc-miR-133a-3p modulates protein expression during JEV infection. Proteomic analysis of cells transfected with the miRNA mimic and infected with JEV revealed a significant downregulation of GRK2, a proviral factor involved in the replication of flaviviruses such as yellow fever virus (YFV), DENV, and HCV (Le Sommer et al., 2012) (Supplementary Fig. 4D). GRK2 was upregulated in NS1-positive 3D4/31 cells, suggesting a potential role in JEV replication. Thus, ssc-miR-133a-3p may exert antiviral effects by binding to the 3′UTR and suppressing GRK2 expression. Moreover, ssc-miR-133a-3p is known to induce cell cycle arrest (Yin et al., 2019), and flaviviruses modulate cell cycle regulation and GPCR signaling pathway for replication (Chan et al., 2018, Valencia et al., 2021, Albarnaz et al., 2014). Gene Set Enrichment Analysis (GSEA) of DEPs in ssc-miR-133a-3p-transfected and infected cells vs. infected cells revealed negative enrichment of pathways related to GPCR signaling, cytokinesis, and microtubule-based movement (Fig. 5H, Supplementary Fig. 4E). This suggests that miR-133a-3p induces cell cycle arrest and modulates GPCR signaling to inhibit JEV replication.

In conclusion, ssc-miR-133a-3p demonstrates a multifaceted antiviral mechanism in JEV infection by targeting the 3′UTR, downregulating GRK2, and modulating critical cellular pathways, ultimately suppressing viral replication and translation.

### Inhibiting hsa-miR-3614-5p increases the virus replication in human macrophages by suppressing the expression of interferon-stimulated genes (ISGs)

Downregulation of ADAR1 leads to the accumulation of unedited double-stranded RNA (dsRNA), triggering a type-I interferon (IFN-I) response that restricts viral replication (de Chassey et al., 2013, Schoggins et al., 2011). This response further induces miR-3614-5p, which modulates the host’s antiviral defense (Diosa-Toro et al., 2017). In this study, proteomic analysis revealed the downregulation of ADARB1, which correlated with the upregulation of miR-3614-5p in THP-1 cells.To confirm the antiviral role of miR-3614-5p during JEV infection, THP-1 cells were transfected with an miRNA inhibitor, and viral gene expression was assessed (Fig. 6A). Inhibition of miR-3614-5p was confirmed by qRT-PCR, which showed reduced expression compared to controls (Supplementary Fig. 5A). This inhibition led to increased NS3 gene expression (Fig. 6B), corroborated by elevated NS1 protein levels (Fig. 6C & 6D). Additionally, plaque assays revealed higher viral titers in miRNA-inhibited cells (Fig. 6E), establishing miR-3614-5p as an antiviral factor suppressing JEV replication.

**Figure 6.**
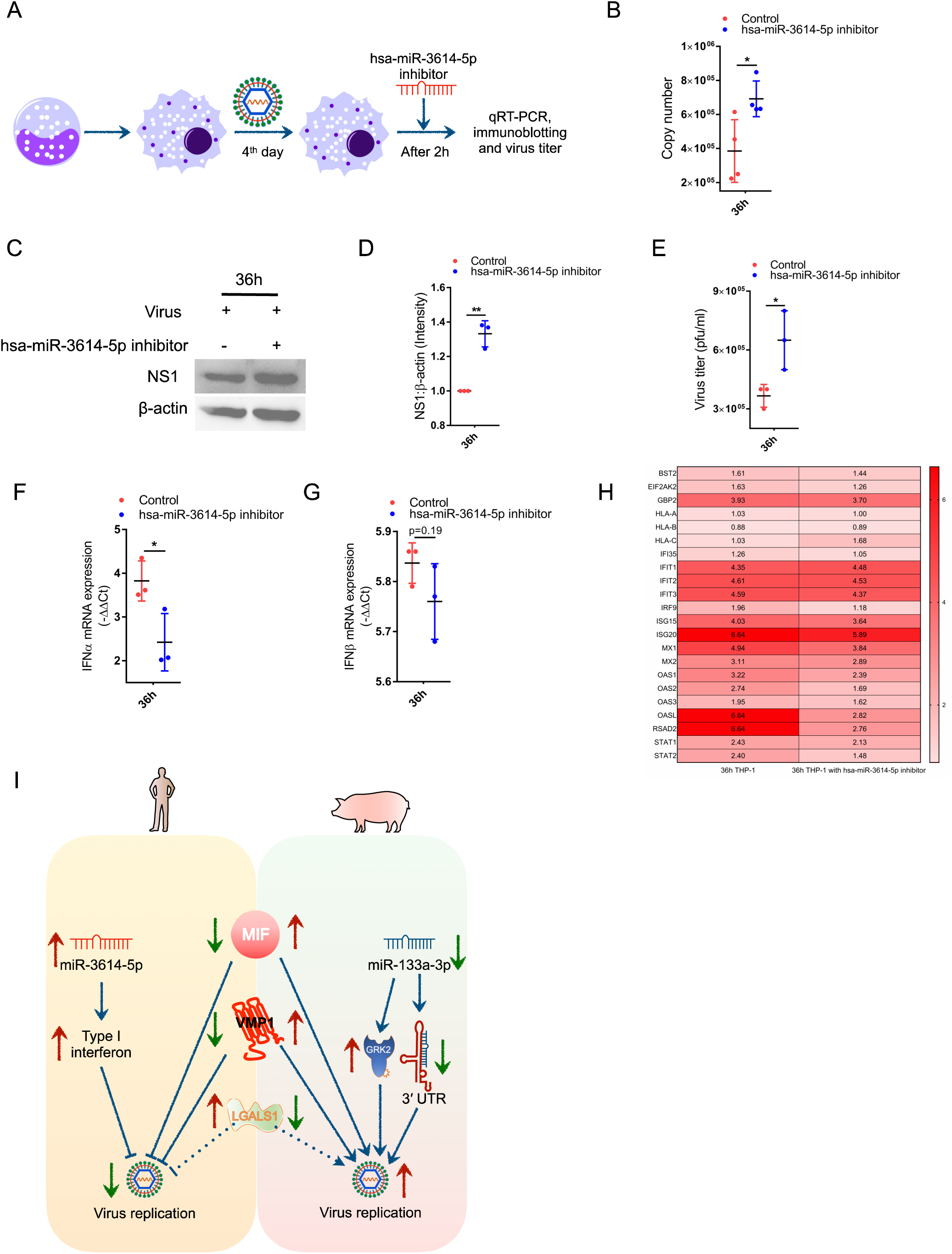
hsa-miR-3614-5p is an antiviral. (A) Workflow showing the protocol for miRNA inhibitor transfection in THP-1 cells. (B) qRT-PCR of the NS3 gene of JEV at 36hpi in THP-1 and inhibitor-transfected THP-1 cells, and the copy number of NS3 was plotted in the y-axis, representing the differences between THP-1 and inhibitor-transfected THP-1. The data represented here were from two independent biological replicates for control, with two biological replicates being repeated one more time and three for inhibitor-transfected cells, with one biological replicate being repeated one more time. Error bars represent the mean with SD. Unpaired t-test. ∗p < 0.05, ∗∗p < 0.01, ∗∗∗p < 0.001, and ∗∗∗∗p < 0.0001. (C & D) Immunoblotting of NS1 protein at THP-1 and inhibitor-transfected THP-1 cells and intensity plot of the ratio of NS1 to β-actin of 36hpi in THP-1 and inhibitor-transfected THP-1. The data represented here were from three independent biological replicates. Error bars represent the mean with SD. Unpaired t-test. ∗p < 0.05, ∗∗p < 0.01, ∗∗∗p < 0.001, and ∗∗∗∗p < 0.0001. (E) Plaque assay to determine the virus titer compared between THP-1 and inhibitor-transfected THP-1 cells at 36hpi. The data represented here were from three independent biological replicates. Error bars represent the mean with SD. Unpaired t-test. ∗p < 0.05, ∗∗p < 0.01, ∗∗∗p < 0.001, and ∗∗∗∗p < 0.0001. (F & G) mRNA expression level of IFNIZI (F) and IFNβ (G) between infected THP-1 vs. control THP-1 and inhibitor-transfected and infected THP-1 vs inhibitor-transfected control THP-1. The data represented here were from three independent biological replicates. Error bars represent the mean with SD. Unpaired t-test. ∗p < 0.05, ∗∗p < 0.01, ∗∗∗p < 0.001, and ∗∗∗∗p < 0.0001. (H) Heatmap showing the abundance ratio of type I interferon signaling pathway-related DEPs between inhibitor-transfected and infected cells vs. infected cells. (I) Graphical illustration of the selected and validated molecular determinants for the role of JEV replication.

Further, to explore the impact of miR-3614-5p on IFN-I signaling, we measured IFN-α and IFN-β levels following miRNA inhibition. IFN-α expression was significantly reduced at the mRNA level, while IFN-β remained unchanged (Fig. 6F & 6G). IFN-α expression was significantly reduced at the mRNA level, while IFN-β remained unchanged (Fig. 6F & 6G). Further, the proteome of miRNA-inhibited cells showed a global reduction in antiviral proteins, including STAT1, STAT2, IRF9, OASs, ISG15, and ISG20, compared to untreated infected samples (Fig. 6H, Supplementary Fig. 5B). This aligns with Vuillier et al. (2022), who reported increased RIG-I and MDA5 expression in HeLa cells following miR-3614-5p mimic transfection. Gene Set Enrichment Analysis (GSEA) further showed negative enrichment of immune response pathways, indicating reduced JAK/STAT signaling activity (Supplementary Fig. 5C). Further, Immunoblotting revealed decreased phosphorylated STAT2 levels following miRNA inhibition (Supplementary Fig. 5D & 5E), supporting the hypothesis that miR-3614-5p regulates JAK/STAT signaling during JEV infection. These findings suggest that miR-3614-5p sustains an antiviral state in JEV-infected THP-1 cells by modulating type-I interferon signaling and restricting viral replication.

## Discussion

Japanese Encephalitis Virus (JEV) transmission to humans leads to severe outcomes, particularly in young children. However, humans act as dead-end hosts in the transmission cycle, as they exhibit lower viremia levels compared to pigs, the amplifying hosts. This pattern is linked to the initial replication of JEV at local lymph nodes, keratinocytes, monocytes/macrophages, and T-lymphocytes before its dissemination to the central nervous system (CNS). To investigate the differences in JEV susceptibility between species, we studied infection in THP-1 macrophages (human) and 3D4/31 macrophages (pig). Interestingly, infection rates were significantly higher in pig macrophages (∼65%) compared to human macrophages (∼18%) despite a lower multiplicity of infection (MOI) for the former. These findings are consistent with García-Nicolás et al. (2019), who observed similar patterns in monocyte-derived dendritic cells.

Multi-omics analyses of NS1-sorted cells revealed 24 potential determinants influencing viral replication and dissemination, including proviral factors like macrophage inhibitory factor (MIF) and vacuole membrane protein 1 (VMP1) and antiviral factors such as galectin-1 (LGALS1), hsa-miR-3614-5p, and ssc-miR-133a-3p. Functional assays confirmed their differential roles in modulating host-specific JEV responses. MIF, known for its role in immune regulation, cell survival, and proliferation (Alampour-Rajabi et al., 2015, Schwartz et al., 2009, Shi et al., 2006), was upregulated in 3D4/31 cells but downregulated in THP-1 cells. Its inhibition in pig macrophages reduced viral load, aligning with studies in DENV infection where MIF deficiency attenuated viral replication and inflammation (Assunção-Miranda et al., 2010). In contrast, exogenous MIF in THP-1 cells enhanced viral replication and protein translation, underscoring its proviral role in JEV infection. Similarly, VMP1, a regulator of autophagy and lipid metabolism (Morishita et al., 2019, Ropolo et al., 2007, Zhao et al., 2017), displayed species-specific expression patterns, with upregulation in 3D4/31 cells and downregulation in THP-1 cells. Knockdown experiments confirmed its critical role in supporting JEV replication and protein translation in pig macrophages. Similarly, the expression of VMP1 supports virus replication and translation in human macrophages.

LGALS1 (galectin-1), an antiviral protein known to attenuate virus adsorption and internalization (Toledo et al., 2014), was significantly downregulated in pig macrophages but marginally upregulated in THP-1 cells. Overexpression of LGALS1 in pig macrophages reduced viral replication without affecting protein translation, suggesting an indirect regulatory role. Interestingly, its inhibition in THP-1 cells increased viral replication without enhancing viral release, hinting at its potential role in modulating viral kinetics rather than egress.

Among the antiviral miRNAs studied, ssc-miR-133a-3p interacted with the 3’UTR of JEV RNA, suppressing replication by interfering with RNA synthesis. Overexpression of ssc-miR-133a-3p in pig macrophages reduced viral replication and GRK2 expression, a key player in flavivirus RNA synthesis. GSEA analysis revealed the downregulation of GPCR signaling pathways, supporting its role as an antiviral determinant. Similarly, hsa-miR-3614-5p, an interferon-inducible miRNA (Vuillier et al., 2022), was upregulated in THP-1 cells during infection and inhibited viral replication by maintaining immune homeostasis. Its inhibition reduced IFN-α expression and ISG levels, leading to increased viral replication and diminished immune response, as confirmed by proteomics and GSEA. In brief, after assessing their roles in different stages of viral kinetics, we confirmed that the upregulation of MIF and VMP1 and downregulation of ssc-miR-133a-3p could make JEV more replicative in pig macrophages (Fig. 6I).

### Conclusion

This study highlights the intricate interplay of host-specific factors in regulating JEV replication and immune responses, identifying key determinants that influence viremia levels and host-specific outcomes. Proviral elements such as MIF and VMP1 promote viral propagation, while antiviral factors like LGALS1, hsa-miR-3614-5p, and ssc-miR-133a-3p restrict viral replication through diverse mechanisms. These insights into host-pathogen interactions provide valuable targets for therapeutic intervention in JEV infection. By elucidating the roles of these factors, the study underscores how pigs serve as amplifier hosts with high viremia levels, while humans, with significantly lower viremia, act as dead-end hosts. Though conducted in vitro using macrophages as a model, this comprehensive research lays the groundwork for further exploration of the molecular determinants of viremia and the development of efficient antiviral strategies to mitigate JEV pathogenesis.

## Materials and Methods

### Cells and virus

Human monocytic cell line: THP-1 (ATCC-TIB-202) and pig alveolar macrophage cell line: 3D4/31 (ATCC-CRL-2844) were maintained in RPMI-1640 with 10% (v/v) Foetal Bovine Serum (FBS) with 1X Gentamicin/Amphotericin Solution. THP-1 and 3D4/31 cell lines were supplemented with 1mM glutamine and 1mM non-essential amino acid (NEAA), and in addition to that, 3D4/31 was supplemented with 1mM sodium pyruvate. Pig Kidney cell line: PS and Human embryonic kidney cell line: HEK293 were maintained in Dulbecco’s Modified Eagle’s Medium (DMEM) with 10% (v/v) FBS with 1X Gentamicin/Amphotericin Solution. The stable cell lines were maintained in their respective media along with puromycin (1ug/ml for THP-1 and 5ug/ml for 3D4/31). All cultures were maintained in the humidified CO_2_ incubator at 37°C with 5% CO_2_. The cells were kept in media along with Plasmocin Prophylactic (Invivogen) to keep them free from mycoplasma.

The infection study was done with a live attenuated vaccine strain of JEV (SA14-14-2). The virus was propagated in the PS cell line, and titer was obtained using plaque assay. Before infecting THP-1, THP-1 was induced into macrophages using 5ng/ml of Phorbol 12-myristate 13-acetate (PMA) and incubated for 3 days, followed by one-day incubation in 10% growth medium without PMA. Virus at multiplicity of infection (MOI) of 7 was used to infect THP-1. For 3D4/31 cells, the cells were seeded for 3 days to attain 70 to 80% confluency. On the third day, the virus at MOI of 1 was added for infection in 2% media. The cells were collected at respective time points for different assays, either at 24hpi or 36hpi or 48hpi.

### Plaque assay

The PS cell line was infected with the virus and kept for 3 days for propagation of the virus. The supernatant was aliquoted and kept at −80°C. To check the titer of the virus stock, the virus was diluted at different dilutions and overlaid in the PS cell line for 2h in 2% DMEM medium, followed by an agarose overlay. The plaque was counted after 5 days and infection for further assay was done according to the virus titer. The same protocol was followed to measure the virus titer after treatment or transfection followed by infection. Briefly, after collecting the cell culture supernatant, the supernatant was centrifuged at 1000 x *g* for 5min at 4 °C and collected. This cleared supernatant was used for plaque assay.

### Plasmids

pLex MCS mCherry Myc (Addgene #128051), pMDLg/pRRE (Addgene #12251), pRSV-Rev (Addgene #12253), pMD2.G (Addgene #12259), pmirGLO, and lentiCRISPR v2 (Addgene #52961) were used in this study. LGASLS1 gene was cloned into pLex MCS mCherry Myc between BamHI and SpeI site, and 3’UTR of JEV was cloned into pmirGLO between SacI and XhoI site. The sgRNAs were cloned in lentiCRISPR v2 at BsmBI site.

### Cloning

The PCR amplicon was generated using Phusion™ High-Fidelity DNA Polymerases (2 U/μL) with standard protocol. LGALS1 gene (CDS) was cloned into pLex MCS mCherry Myc between BamHI and SpeI site, and 3’UTR of JEV was cloned into pmirGLO between SacI and XhoI site. The sgRNAs (100µM each) were annealed along with phosphorylation of 5′ end using T4 PNK. The annealed oligos were used for ligation in plasmid at a concentration of 500nM. Then, the sgRNAs against MIF were cloned into lentiCRISPR v2 using BsmBI restriction enzyme. VMP1 gene was cloned into pIRES2-EGFP using BamHI and XhoI enzymes. For the ligation of the insert and vector, T4 ligase was used. The transformation of the ligated product was done in Mach1 T1^R^ competent cells. The positive colony was selected using colony PCR and double RE digestion, and the positive colony was cultured in large volumes for further transfection experiments. The details about the sequence used for the cloning experiment are mentioned in Supplementary Table 1.

### Immunoblotting

The cells were washed at the mentioned time points, put in M-PER reagent with 1x protease inhibitor cocktail for lysis, and kept at −80 °C. Then, the cells were centrifuged at 15000 rpm for 20min at 4 °C. The protein was quantified using the BCA protein assay kit, and 8 to 10ug proteins were resolved in 12% SDS-PAGE gels and transferred onto a polyvinylidene difluoride (PVDF) membrane, followed by blocking for 2h. Then, the membrane was incubated with the primary antibody and followed by the secondary antibody. The washing was done with 1x TBS with 0.1% Tween 20 between the incubation steps. The blot was developed against GAPDH or β-Actin after stripping and reprobing with respective antibody. The immunoblotting was done with antibodies: anti-NS1 (1:10000), anti-NS3 (1:10000), anti-MIF (1:2000), anti-LGALS1 (1:5000), anti-Phospho-Stat2 (Tyr690) (1:1000), anti-GAPDH (1:1000), anti-β-Actin (1:1000), HRP-conjugated goat anti-mouse IgG (1:15000) and HRP-conjugated goat anti-rabbit IgG antibody (1:5000) were used. The blot was developed using SuperSignal West Pico or femtoLUCENT PLUS HRP Chemiluminescent Substrate, and the image was captured in ChemiDoc (Biorad, USA) or iBright 1500 (Thermo Fisher Scientific Inc., USA).

### Flow cytometry analysis and sorting of NS1-positive cells

Trypsinized cells were pelleted down and fixed with 4% paraformaldehyde for 10 min at 4 °C. Fixed cells were blocked with the blocking buffer (3% BSA with 0.1% Triton-X-100 in PBS) for 1h. After blocking, cells were incubated with anti-NS1 (1:500) for 1h, followed by APC-conjugated anti-mouse IgG antibody (1:1000) for 30min. The antibody incubation was done in dilution buffer (1% BSA with 0.1% Triton-X-100 in PBS), and in between, washing was done with washing buffer (1% BSA with 0.1% Tween-20 in PBS). Then, the cells were resuspended in the washing buffer and analyzed using BD FACSMelody™ Cell Sorter. The data was analyzed using FlowJo v. 10. For sorting of NS1 positive cells, the gating strategy was employed with respect to control cells. Sorted cells were later centrifuged and processed for proteomics and miRNomics.

### Gene expression analysis by qRT-PCR

After infection, the cells were harvested, and RNA was isolated using Qiagen RNeasy mini kit or NucleoSpin RNA Plus kit. RNA was reverse transcribed using PrimeScript 1^st^ strand cDNA Synthesis Kit. For quantitative Real-time PCR (qRT-PCR) using the primers (Supplementary Table 2), SYBR Green PCR-Master mix was used with cycling conditions: initial denaturation at 95°C for 1 min, followed by 36 cycles of 95°C for 10 s and 62.2°C for 30 s, using CFX96 Touch Real-Time PCR Detection System (Bio-Rad). The standard curve for the NS3 gene was used to determine the copy number using the same primer mentioned in Supplementary Table 2. Except for NS3, IFNIZI, IFNβ, and GAPDH gene, the annealing temperature for VMP1 and β-actin was set to 60 °C. Relative expression was determined using the −ΔΔCt method after normalizing the Cq value with the Cq value of glyceraldehyde-3-phosphate dehydrogenase (GAPDH).

RNA was isolated using RNAiso Plus reagent for miRNA quantification, followed by DNase I treatment. The RNA (375 ng) was converted into cDNA using the Mir-X™ miRNA First Strand Synthesis Kit. The qRT-PCR was done using the earlier protocols with slight modification by considering annealing temperature at 60°C, and for normalization, expression of U6 gene was used.

The NS3 amplicon was amplified using the primer mentioned in Supplementary Table 2. The amplicon on serial dilution ran for qRT-PCR. The Cq value was plotted in correspondence to its molecule number to make a standard curve. The generated standard curve resulted in the following equation-Y= −3.563X + 38.245, where Y represents the Cq value and X represents log10. A high coefficient of determination (R^2^ = 0.999) indicated the reliability of the standard curve. Viral copy numbers were computed and are presented as copy number per 10 ng total RNA.

### Enzyme**-**linked immunosorbent assay (ELISA)

The supernatants from THP-1 cells (control and infected) were harvested after 24h, 36h, and 48h. The cytokines IL-1β, IL-6, and TNF-α were measured using sandwich ELISA kits (R&D Systems) per the standard protocol. The absolute concentrations were estimated using a standard curve and expressed as picogram per milliliter.

### Generation of stable cell line

A third-generation lentiviral system was used to generate stable cell lines. The coding sequence (CDS) of LGALS1 was retrieved from cDNA using the primer mentioned in Supplementary Table No. 1 and was cloned before mCherry gene to make fused LGALS1-mCheery into pLex MCS mcherry myc plasmid. HEK293 cells were seeded and kept overnight before transfection. The two packaging plasmids-pMDLgpRE and pRSVRev with LGALS1 containing plex mcs mcherry plasmid and pMD2.g were transfected into HEK293 cells with Lipofectamine 3000. The supernatant was collected after 48h, filtered, and stored at −80°C. The viral transduction was done to the 3D4/31 cells using 8ug/ml polybrene. After 2 days, the cells were kept for selection in 5ug/ml puromycin. The LGALS1 expressing stable 3D4/31 cells were used for further experiments. For generating stable MIF KO 3D4/31 cells, instead of plex mcs mcherry plasmid, lentiCRISPR v2 containing sgRNA was transfected in HEK293. As in this study, two individual guide RNAs were used, we mixed the supernatant from HEK293 after transfection in 1:1 ratio before transduction.

### Transfection of plasmid

THP-1 cells were seeded and infected using the methods mentioned above. After 2h of infection, pIRES2-EGFP and pIRES2-EGFP-VMP1 plasmids were transfected using Lipofectamine 3000 as per the standard protocols, and after 6h, fresh media was replenished. Further assay was performed at respective time points.

### Gene silencing using siRNA

The 3D4/31 cells were seeded, and siNC (25nM) or siVMP1(25nM) were transfected using Lipofectamine 3000 before 24h of infection (Supplementary Table 3). Two siRNAs against VMP1 were mixed for transfection. After 6h of transfection, the cells were washed and put in fresh media.

### Transfection of miRNA mimic or miRNA inhibitors

After infection, followed by washing, the ssc-miR-133a-3p mimic (50nM) or hsa-miR-3614-5p miRNA inhibitor (25nM) was transfected using Lipofectamine 3000 (Supplementary Table 3). After 6h, the cells were washed and put in the growth medium for respective time points.

### Drug treatment for inhibiting MIF and LGALS1

Known MIF inhibitor: ISO-1 was given at a concentration of 20uM. DMSO or ISO-1 was added 12h before infection and 24h after infection in 3D4/31 cells. LGALS1 inhibitor-OTX-008 at a concentration of 30uM or ethanol was added 48h before infection in PMA-induced THP-1 cells.

### Human recombinant MIF

Recombinant MIF at 100ng/ml or 0.1% BSA was added in THP-1 cells 2h after overlaying the virus, and the cells were collected for respective time points.

### *In silico* analysis of binding of 3**′**UTR and ssc-miR-133a-3p

The 3′UTR of JEV (SA14-14-2: MK585066.1) was retrieved from NCBI. The ssc-miR-133a-3p sequence was retrieved from miRbase. BiBiServ was used to predict the *in-silico* binding energy between the target and miRNA.

### Dual luciferase assay

HEK293 cells were seeded overnight. The next day, pmirGLO or pmirGLO-3′UTR was transfected alone or with ssc-miR-133a-3p mimic in HEK293 cells with Lipofectamine 3000. After 48h of transfection, the cells were washed with 1X PBS and put in 100ul of PLB buffer. The plate was kept in the shaker for 15 minutes, and later, the cells in the buffer were collected and stored at −80°C. The samples were prepared, and luminescence was taken per the standard protocol in the Dual-Luciferase Reporter Assay System. The normalization was done with the luminescence of the Renilla luciferase gene.

### Sample preparation for LC-MS/MS of NS1-sorted cells

The sorted infected and control cells were washed with PBS and centrifuged. The pellet was resuspended in 50 mM Tris buffer (pH 8.0) with 2% Sodium deoxycholate. The samples were incubated at 98°C for 30min followed by 60°C for 2h. Then, the samples were stored at −80°C for further processing. The cell lysates were sonicated and centrifuged. After quantifying the protein concentration using the BCA protein assay kit, Dithiothreitol (DTT) was added and incubated at 56°C for 1h to reduce the disulfide bond. To further inhibit the bond reformation, Iodoacetamide (IAA) was added and kept at room temperature in the dark for 1h. Trypsin/Lys-C Mix was added at a final protease: protein ratio of 1:20 (w/w) and incubated at 37 °C overnight. Then, 2.5% formic acid was added to stop enzymatic lysis. Then, the samples were filtered using protein concentrators, 10K MWCO, followed by C18 column purification. The purified peptides were vacuum-dried and stored at −80°C before running in LCMS/MS. To check the global proteome spectrum after mimic or miRNA inhibitor transfection, we collected the cells at 36hpi as whole cell lysate and processed them as per the above-mentioned protocols, but instead of Trypsin/Lys-C Mix, only Trypsin was used for protein digestion.

### LC-MS/MS analysis

The vacuum-dried peptides were reconstituted in 0.3% formic acid and injected into LC-MS/MS (Q Exactive, Thermo Scientific). The running time for each sample was 240min. The MS scan had a range of 375 to 1600 m/z and a resolution of 60000. The dd-MS scan had a resolution of 15,000.

### Analysis of proteomics data

The raw data was analyzed using Proteome Discoverer (v2.4.1.14). The reference databases were *Homo sapiens* for samples from THP-1 and *Sus Scrofa* for samples from 3D4/31. FDR was < 0.01 for selecting proteins with high confidence. Label-free quantification was based on the ratio of the abundance of infected samples to the abundance of control samples. The proteins were considered differentially expressed proteins (DEPs) if the abundance ratio had an adjusted p-value < 0.05. The PCA was generated using FactoMineR, and the volcano plot and dot plot were created using ggplot2 package in RStudio.

### Gene Ontology analysis

The upregulated and downregulated DEPs of both time points were analyzed for their association with enriched pathways using ClueGO (Bindea et al., 2009). The biological processes were considered for DEPs of sorted samples. The pathways or processes were considered significant if the p-value corrected with Bonferroni step-down was < 0.05.

### Gene Set Enrichment Analysis of proteomics data

The protein names of pigs were matched with human orthologs and analyzed the association of DEPs in enriched biological processes of humans by GSEA (Subramanian et al., 2005) using the pre-ranked file, and the cut-off was set for FDR <0.05. The DEPs of humans were analyzed for enriched biological processes using GSEA using the pre-ranked file, and the cut-off for selection of biological processes was considered as p-value < 0.05.

### Sample preparation for miRNA sequencing

Total RNA from sorted cells was isolated using the standard manufacturer’s protocol using TRIzol. Following the manufacturer’s protocol, the library was prepared using TruSeq Small RNA Library Preparation Kits (illumina, USA). A hundred nanograms of total RNA was taken for library preparation. The quality of the libraries was measured using the TapeStation system (Agilent, USA). Libraries were quantified using a Qubit 2.0 Fluorometer (Life Technologies). The high-throughput sequencing was performed on Illumina – NovaSeq 6000 (illumina, USA) (50 bp paired-end) per the manufacturer’s protocol.

### Analysis of miRNomics data

Known mature miRNAs and precursor sequences for humans and pigs were retrieved from the miRBase database. The mapping and quantification of miRNAs were done using the miRDeep2 tool (Friedländer et al., 2012). After filtering the reads (length ≥18 nt) and adapter trimming, the filtered reads were mapped to the indexed human or pig genome using the mapper module. After mapping, the aligned reads were quantified using the quantifier.pl module. The differentially expressed miRNAs (DEmiRNAs) were identified using DESeq2, and the DEmiRNAs were considered if the padj was <0.05 for downstream analysis. The PCA and volcano plot were generated using FactoMineR and ggplot2 packages, respectively, in RStudio.

### Screening strategy for selecting the factors

The screening strategy involved four steps: (1) Identifying DEPs linked to viral processes using g:Profiler, (2) Cross-referencing DEPs with previously reported RNAi, CRISPR/Cas9, or immunoprecipitation studies to determine proviral or antiviral host factors (Savidis et al., 2016, Zanini et al., 2018, Krishnan et al., 2008, Hoffmann et al., 2018, Zhang et al., 2016, Aviner et al., 2021, Shah et al., 2018, Kovanich et al., 2019), (3) Prioritizing DEPs based on different expression, and (4) Screening DEPs associated with metabolic pathways. A literature-driven approach was also applied to DEmiRNAs to identify key regulatory determinants in flavivirus infections.

### Quantification and statistical analysis

Generally, all the experiments were done for three independent biological replicates unless otherwise stated. The densitometry analysis for immunoblot was done using Image Lab software (BioRad) or iBright Analysis software (Thermo Scientific). The graphs were prepared using GraphPad prism 7, and the statistical significance was calculated using unpaired Student’s t-test or two-way ANOVA (mentioned in the figure legend).

## Data Availability Statement

The proteome datasets of this study were submitted to ProteomeXchange Consortium via the PRIDE partner repository with the dataset identifiers PXD060527 and PXD060545, and miRNome data were submitted to NCBI GEO with accession number GSE289291.

## Acknowledgments

We thank the Department of Biotechnology, India, for providing financial support for conducting research. MRP and TA acknowledge CSIR, India, for providing fellowship. We thank Dr. Nagendra R. Hegde for assisting in analysis. We also thank Dilna Vasanthakumari and Nilanjana Ganguli for assisting with LC-MS/MS studies and Rama Devi and Kreethi for assisting in flow cytometry analysis.

## Authorship contribution statement

RKG and MRP conceptualized and designed the study. MRP and TA performed experiments. MRP performed formal analysis. BD, RKG, BRG, SSM, and GTS supplied resources. RKG, BD, GTS, and SSM handled funding acquisition and project administration. MRP did the initial drafting, and SK, BD and RKG did the reviewing and editing. All authors read and approved the manuscript.

## Supplementary figure legends

**Supplementary Figure 1.** Dysregulated proteins and miRNAs and their associated pathways of 3D4/31.

(A) PCA plot showing the demarked clustering of control and infected samples at 36hpi. (B). Volcano plot rendering the downregulated (marked as green), upregulated (marked as red), and nonsignificant (marked as grey) DEPs at 36hpi.

(C) Dot plot illustrating the enriched biological processes for the upregulated DEPs at 36hpi. (D) Dot plot depicting the enriched pathways for downregulated DEPs at 36hpi. (E) PCA plot showing the demarcated clustering of control and infected samples at 48hpi. (F). Volcano plot rendering the downregulated (marked as green), upregulated (marked as red), and nonsignificant (marked as grey) DEPs at 48hpi.

(G) Dot plot illustrating the enriched biological processes for the upregulated DEPs at 48hpi. (H) Dot plot depicting the enriched pathways for downregulated DEPs at 48hpi. (I) PCA plot showing the demarked clustering of DEmiRNAs of control and infected samples at 36hpi. (J) Volcano plot rendering the downregulated (marked as green), upregulated (marked as red), and nonsignificant (marked as grey) DEmiRNAs at 36hpi.

**Supplementary Figure 2.** Dysregulated proteins and miRNAs and their associated pathways of THP-1.

(A) PCA plot showing the demarked clustering of control and infected samples at 36hpi. (B) Volcano plot rendering the downregulated (marked as green), upregulated (marked as red), and nonsignificant (marked as grey) DEPs at 36hpi. (C) Dot plot illustrating the enriched biological processes for the upregulated DEPs at 36hpi. (D) Dot plot depicting the enriched pathways for downregulated DEPs at 36hpi. (E) PCA plot showing the demarcated clustering of control and infected samples at 48hpi. (F). Volcano plot rendering the downregulated (marked as green), upregulated (marked as red), and nonsignificant (marked as grey) DEPs at 48hpi. (G) Dot plot illustrating the enriched biological processes for the upregulated DEPs at 48hpi. (H) Dot plot depicting the enriched pathways for downregulated DEPs at 48hpi. (I) PCA plot showing the demarked clustering of DEmiRNAs of control and infected samples at 36hpi. (J) Volcano plot rendering the downregulated (marked as green), upregulated (marked as red), and nonsignificant (marked as grey) DEmiRNAs at 36hpi.

**Supplementary Figure 3.** Selection of proviral and antiviral determinants responsible for distinct virus levels in both hosts.

(A) Plan was conducted for selecting the key determinants through global profiling of the proteome and miRNome.

**Supplementary Figure 4.** ssc-miR-133a-3p has antiviral activity.

(A) In silico analysis of binding of the 3’UTR of JEV with ssc-miR-133a-3p. (B) The plot represents the ratio of Fluc to Rluc, showing the differences between control and mimic transfected HEK293 transfected with pmirGLO. The data represented here were from three independent biological replicates. (C) qRT-PCR of ssc-miR-133a-3p expression at 36h in 3D4/31 with or without mimic transfection. The data represented here were from three independent biological replicates. (D) Volcano plot plotted for infected 3D4/31 vs. control 3D4/31 (First one) and for mimic transfected and infected 3D4/31 vs mimic transfected control 3D4/31 (Second one). The apoptotic-related DEPs are marked in the plot. The green color represents the downregulated DEPs, and the red represents the upregulated DEPs. (E) GSEA enrichment plot showing the enrichment score of biological processes using the pre-ranked file between mimic-transfected and cells vs. infected cells.

**Supplementary Figure 5.** hsa-miR-3614-5p has antiviral activity.

(A) qRT-PCR of hsa-miR-3614-5p expression at 36hpi in THP-1 with or without inhibitor transfection. Data represented here were from three independent biological replicates. (B) Volcano plot plotted for infected THP-1 vs. control THP-1 (left one) and for inhibitor-transfected and infected THP-1 vs inhibitor-transfected control THP-1 (right one). The immune response-related DEPs are marked in the plot. The green color represents the downregulated DEPs, and the red represents the upregulated DEPs. (C) GSEA enrichment plot showing the enrichment score of biological processes using the pre-ranked file between inhibitor-transfected and infected cells vs. infected cells. (D & E) Immunoblotting of pSTAT2 protein at THP-1 and inhibitor-transfected THP-1 cells and intensity plot of the ratio of pSTAT2 to β-actin of 36hpi in THP-1 and inhibitor-transfected THP-1. The data represented here were from three independent biological replicates. Error bars represent the mean with SD. Unpaired t-test. ∗p < 0.05, ∗∗p < 0.01, ∗∗∗p < 0.001, and ∗∗∗∗p < 0.0001.

