## Supplementary File 1 for "Multi-omics analysis uncovers differential growth kinetics of Japanese Encephalitis Virus in human and porcine macrophages"

**Supplementary Discussion**

C1QBP or gC1qR localized in mitochondria moves to the cytoplasm upon infection (Xu et al., 2009) to regulate innate and adaptive immunity. HCV core protein interacts with gC1qR to inhibit the proliferation of T-cells from making persistent infections (Kittlesen et al., 2000). Upregulation of this scuttles RIG-I and MDA5 signaling in Sendai virus infection, aiding the viral replication (Xu et al., 2009). The positive relationship of C1QBP with viral replication is also observed in respiratory syncytial virus (RSV) (Hu et al., 2017). Conversely, this downregulation in THP-1 cells may augment antiviral molecules and signaling pathways to suppress viral replication.

CHK2, a protein kinase, is induced in response to DNA damage and regulates cell cycle arrest (Hirao et al., 2000). G2/M phase arresting, activating dsDNA damage response pathway in response to double-strand breaks, and mutation in cellular genes are induced by HCV infection (Wu et al., 2008, Machida et al., 2004). Attenuating ATM or CHK2 repressed the HCV replication (Ariumi et al., 2008). Likewise, there was a report of persistent infection in neural progenitor cells in JEV infection by manipulating the cell cycle (Das & Basu, 2008). Chan et al. (2018) reported that hijacking CHEK2 by JEV to prolong the cell cycle arrest has been advantageous for virus replication. Nevertheless, decreasing this expression in THP-1 cells can negatively modulate viral replication.

Silencing of COIL, an essential component of the Cajal body (CB), reduces the expression of West Nile virus (WNV) and DENV in HeLa cells (Krishnan et al., 2008). Even though there were reports of localization of the NS5 gene in CB of ZIKV (Coyaud et al., 2018, Grant et al., 2016, Kesari et al., 2020), there is no full proof report on its proviral activity. The report of Coyaud et al. (2018) presented the speculations that the interaction may either impair the processing of host mRNAs to promote viral RNA translation by degrading CB or hijack the CB for the maturation of vRNA. The role of upregulated COIL in 3D4/31 may imply some insights into JEV replication or translation.

Succinate exerts an antiviral effect on IAV (Guillon et al., 2022), ZIKV, and WNV (Daniels et al., 2019). In the Influenza A virus, succinate promotes the succinylation of nucleoprotein and its retention (Guillon et al., 2022). Blocking Succinate dehydrogenase (SDH) generates an antiviral state by piling up succinate in the cells (Daniels et al., 2019). As oxidative phosphorylation and Kreb's cycle play a pivotal role in virus pathogenesis, succinate accumulation, and fumarate depletion may worsen the condition for the virus for its replication. Upregulation of SDHC and SDHD, components of SDH in 3D4/31 cells at 36h, may contribute positively to viral replication by converting succinate to fumarate.

Prolyl hydroxylation is a cotranslational modification responsible for folding Flavivirus: DENV and ZIKV polyproteins. A decrease in prolyl hydroxylation aggregates the NS2B and degrades other viral proteins. This minimizes the viral titer (Aviner et al., 2021). The prolyl hydroxylation was done by the ER-resident enzyme- prolyl hydroxylase viz. P4HA1 and P4HA2, and P3H1 and P3H2. The upregulation of P4HA2 in 3D4/31 cells at 48h may protect the viral proteins from aggregation, mainly NS2B.

Signal peptidase complex subunit 1 (SPCS1), a microsomal signal peptidase complex component, is essential to cleave flavivirus structural proteins and their secretions. Curtailing SPCS1 expression reduced the flavivirus yield (Zhang et al., 2016). In flavivirus-like HCV infection, it interacts with E2 and NS2 (Suzuki et al., 2013); in JEV infection, it interacts with NS2B (Ma et al., 2018). This interaction may regulate the virus's life cycle by assisting in virus assembly. Upregulation of SPCS1 may assist in viral assembly and promote viral replication in 3D4/31 at 36h. However, SPCS2, which is proviral in DENV and WNV infection (Zhang et al., 2016, Zanini et al., 2018), was downregulated in 3D4/31 at 48h. This may be because the infectivity at 48h was reduced compared to 36h in 3D4/31 cells.

G protein-coupled receptor kinases (GRKs) phosphorylate G protein-coupled receptors (GPCRs) (Pitcher et al., 1998). Among GRKs, one important GRK-GRK2 plays canonical and non-canonical activity by interacting with GPCR and other non-GPCR membrane receptors and proteins (Penela et al., 2010). This non-canonical interaction exerts a positive modulation in flaviviral replication viz. in yellow fever virus (YFV), DENV, and HCV (Le Sommer et al., 2012) and in other viruses like in Influenza A virus (IAV) (Yángüez et al., 2018). It favors the flaviviral life cycle by supporting both entry (in YFV) and RNA synthesis but not translation (Le Sommer et al., 2012). An increase in GRK2 may foster viral replication, not translation, at 48h in 3D4/31 cells.

Certain proteins involved in mRNA remodeling, decaying, and translational suppression are localized in distinct cytoplasmic domains like processing (P), XRN1, or GW bodies (Eulalio et al., 2007, Eystathioy et al., 2002, Eystathioy et al., 2003). The P-bodies contain decapping proteins, deadenylase complex, exonuclease, and decapping activators- LSM1 to LSM7. LSM1-7 plays a role in the RNAi pathway. Depleting LSM1 in HeLa cells reduces WNV infection by ~70% (Chahar et al., 2013), but the exact mechanism remains unknown. In addition, LSM1 binds to 3' UTR of DENV to increase viral replication (Dong et al., 2015). However, speculation is that P-body components are bridging 3' to 5' termini for efficient replication and/or translation, as a report published by Ward et al. (2011) described the interaction of stress granules components viz. DDX6, G3BP1, G3BP2, Caprin1, and USP10 with the 3' terminus of DENV for efficient replication. The downregulation of LSM1 may not promote viral replication in THP-1 cells at 36hpi.

Polymerase-associated factor 1 complex (PAF1C), composed of PAF1, LEO1, CTR9, RTF1, and CDC73 (Mueller et al., 2002), has been known to regulate gene expression related to antiviral and inflammatory response positively (Parnas et al., 2015). For its negative implication in virus kinetics, it was reported as an antiviral factor for DENV and ZIKV (Shah et al., 2018, Kovanich et al., 2019, Petit et al., 2021). Remarkably, PAF1 positively impacts JEV replication plausibly by interacting with NS5 (Kovanich et al., 2019). As in our study, it was downregulated in THP-1 at 36hpi, which might curtail JEV progression in THP-1.

The membrane trafficking molecule TMED2 is a part of the coatomer complex (COPI) vesicle-mediated retrograde trafficking and trafficking from Golgi to the plasma membrane (Fiedler et al., 1996, Goldberg, 2000). Knocking down and overexpression of TMED2 decrease and increase DENV infection, respectively (Zanini et al., 2018). Upregulation in 3D4/31 cells at 36h may positively affect viral replication.

Cholesterol biogenesis is an essential regulator in viral pathogenesis, starting from entry to evasion. One important molecule involved in cholesterol synthesis is HMG-CoA reductase (HMGCR). HMGCR converts 3-hydroxy-3-methylglutraryl-CoA to Mevalonic acid (Ku, 1996). HMGCR in DENV positively affects viral replication (Rothwell et al., 2009, Mackenzie et al., 2007) by surging cholesterol levels at the replication site. Moreover, the Inhibition of HMGCR by lovastatin reduces DENV production by interfering with viral assembly in PBMCs and other cells (Rothwell et al., 2009, Martínez-Gutierrez et al., 2011, Soto-Acosta et al., 2013, Bryan-Marrugo et al., 2016). Downregulation of HMGCR at 36h in THP-1 cells may inhibit viral replication.

High reactive oxygen species (ROS) production leads to oxidative stress. Oxidants, viz. NO, O_2_.-, OH. and its by-product H_2_O_2_ modulate cellular responses, viral replication, host defenses, and viral pathogenesis (Zhang et al., 2019). After JEV infection, the elevated O_2_.- triggers superoxide dismutase (SOD) for suppressing superoxide (Kumar et al., 2009). JEV infection in microglia activates pro-inflammatory cytokines- IL1β and IL-18, mediated by ROS production (Kaushik et al., 2012). These same proteins were seen as highly expressed in THP1 at 36h. Catalase (CAT), in the peroxisome, converts H_2_O_2_ to H_2_O and O_2_, further limiting ROS production. Moreover, oxidative stress positively modulates flaviviral RNA replication, and antioxidant has a negative impact on viral replication (Gullberg et al., 2015). However, in our study, there was an opposite expression pattern of CAT, upregulated in 3D4/31, and SOD1, downregulated in THP1. This may give another aspect of ROS production to viral replication, which was seen in *Aedes aegypti*; CAT has a positive correlation with viral replication (DENV, not ZIKV) (Oliveira et al., 2017).

The small RNA regulates antiviral responses exerted by viral invasion from the virus or the host. miR-146a, an IFN production regulator, impaired the production of interferon, reported in human monocytes (THP-1 cell) (Wu et al., 2013). In another report from Nahand et al. (2020), it was shown that this miRNA is making different immune cells more susceptible to viral infection by exogenous release. The report proposed this as a biomarker by comparing transcriptomics samples of DENV-infected patients (Xie et al., 2021). Notably, the same report was published concerning JEV, showing that the expression of miR-146a suppresses the NF-kB signaling pathway by inhibiting TRAF6 (Sharma et al., 2015). Moreover, the report from ZIKV corroborates the same (Shukla et al., 2021). Another important miRNA- miR-150, was reported to be a biomarker in DENV infection. Transfection of mir-150 mimic in DENV-infected CD14+ cells suppressed SOCS1 to promote viral replication (Chen et al., 2014). Interestingly, a report showed that the expression of human miR-150a subverts the antiviral effector molecules' activities in mosquitoes (Zhu et al., 2021). They reported that miR-150a hijacks argonaute 1 protein-mediated RNA interference (RNAi) and restricts chymotrypsin expression, a potent antiviral in mosquitoes. Over that, the report from Goswami et al. (2017) shows a negative correlation between TNF-a and miR-150-5p expression in JEV patients. They speculated that it negatively impacts TGF-b signaling, MAPK, and NF-kB signaling. In addition, it also elicits immunotolerance by impairing cytokine production (Sang et al., 2016). Wnt signaling is known to be a proviral signaling pathway molecule (Qin et al., 2016) besides its role in apoptosis, cell proliferation, and development (Cha et al., 2012). Its regulation by miR-34a induces IFN response, impacting flaviviral replication by increasing the binding of GSK3b with TBK, leading to the activation of IRF3 (Smith et al., 2017). Downregulation of miR-34a impairs the IFN response to reverse the inhibitory effect of antiviral responses. Deducing from another study, the upregulation of miR-146a-5p and miR-150 and downregulation of miR-34a may promote viral replication in 3D4/31.

miR-124-3p has a role in interfering with the role of DNM2, a GTPase responsible for vesicle separation, which, on upregulation, inhibits the JEV replication in PK-15 cells (Yang et al., 2016). In ZIKV, the upregulation of miR-124-3p downregulates TFRC, subsequently implying microcephaly phenotype (Dang et al., 2019). Including a single copy of the target site for miR-124 in the chimeric TBEV/DENV genome attenuates virus replication in primary neuronal cells (Heiss et al., 2011). Corroborating from the previous study, it can be hypothesized that the upregulation of hsa-miR-124-3p in THP1 at 36hpi may curtail the virus replication in human macrophages.
