## Supplementary Figure for "Multi-omics analysis uncovers differential growth kinetics of Japanese Encephalitis Virus in human and porcine macrophages"

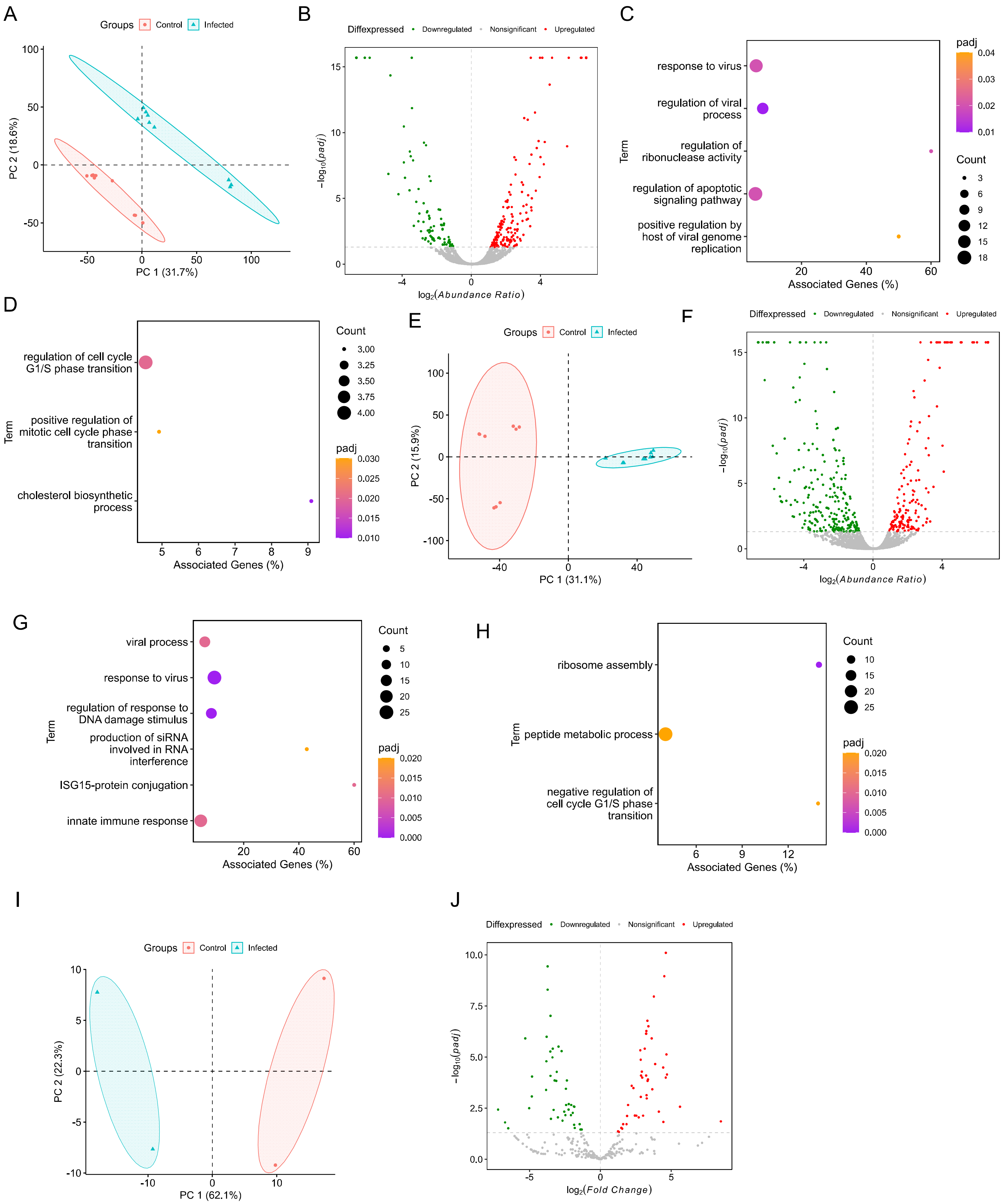

A
Groups
Control
Infected
PC 2 (18.6%)
PC 1 (31.7%)
B
Diffexpressed
Downregulated
Nonsignificant
Upregulated
$-\log_{10}(padj)$
$\log_2(Abundance\ Ratio)$
C
response to virus
regulation of viral process
regulation of ribonuclease activity
regulation of apoptotic signaling pathway
positive regulation by host of viral genome replication
Term
Associated Genes (%)
padj
Count
D
regulation of cell cycle G1/S phase transition
positive regulation of mitotic cell cycle phase transition
cholesterol biosynthetic process
Term
Associated Genes (%)
Count
padj
E
Groups
Control
Infected
PC 2 (15.9%)
PC 1 (31.1%)
F
Diffexpressed
Downregulated
Nonsignificant
Upregulated
$-\log_{10}(padj)$
$\log_2(Abundance\ Ratio)$
G
viral process
response to virus
regulation of response to DNA damage stimulus
production of siRNA involved in RNA interference
ISG15-protein conjugation
innate immune response
Term
Associated Genes (%)
Count
padj
H
ribosome assembly
peptide metabolic process
negative regulation of cell cycle G1/S phase transition
Term
Associated Genes (%)
Count
padj
I
Groups
Control
Infected
PC 2 (22.3%)
PC 1 (62.1%)
J
Diffexpressed
Downregulated
Nonsignificant
Upregulated
$-\log_{10}(padj)$
$\log_2(Fold\ Change)$

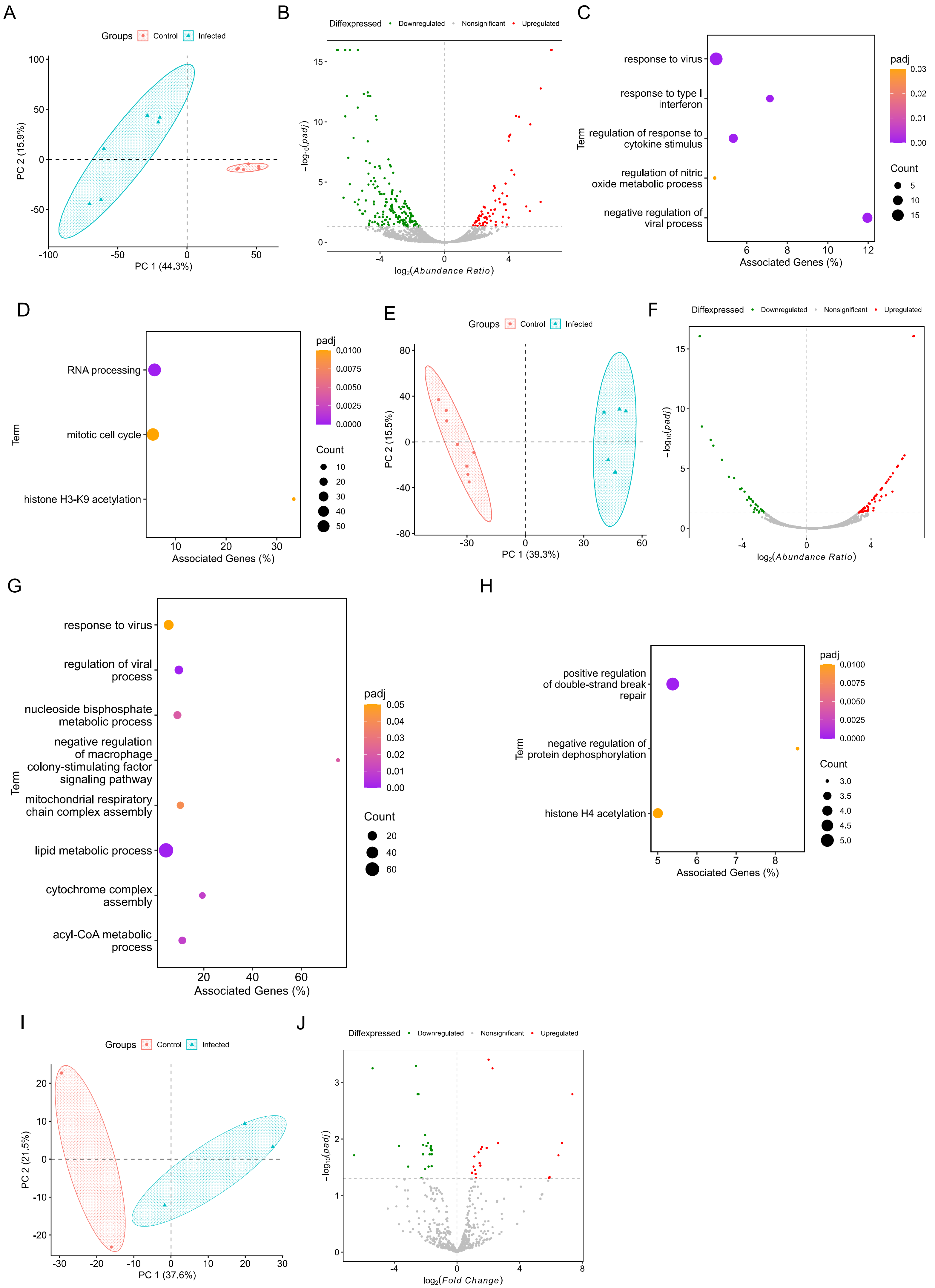

A
Groups Control Infected
PC 1 (44.3%)
PC 2 (15.9%)
B
Diffexpressed Downregulated Nonsignificant Upregulated
log2(Abundance Ratio)
−log10(padj)
C
response to virus
response to type I interferon
regulation of response to cytokine stimulus
regulation of nitric oxide metabolic process
negative regulation of viral process
Term
Associated Genes (%)
padj
Count
D
RNA processing
mitotic cell cycle
histone H3-K9 acetylation
Term
Associated Genes (%)
padj
Count
E
Groups Control Infected
PC 1 (39.3%)
PC 2 (15.5%)
F
Diffexpressed Downregulated Nonsignificant Upregulated
log2(Abundance Ratio)
−log10(padj)
G
response to virus
regulation of viral process
nucleoside bisphosphate metabolic process
negative regulation of macrophage colony-stimulating factor signaling pathway
mitochondrial respiratory chain complex assembly
lipid metabolic process
cytochrome complex assembly
acyl-CoA metabolic process
Term
Associated Genes (%)
padj
Count
H
positive regulation of double-strand break repair
negative regulation of protein dephosphorylation
histone H4 acetylation
Term
Associated Genes (%)
padj
Count
I
Groups Control Infected
PC 1 (37.6%)
PC 2 (21.5%)
J
Diffexpressed Downregulated Nonsignificant Upregulated
log2(Fold Change)
−log10(padj)

A

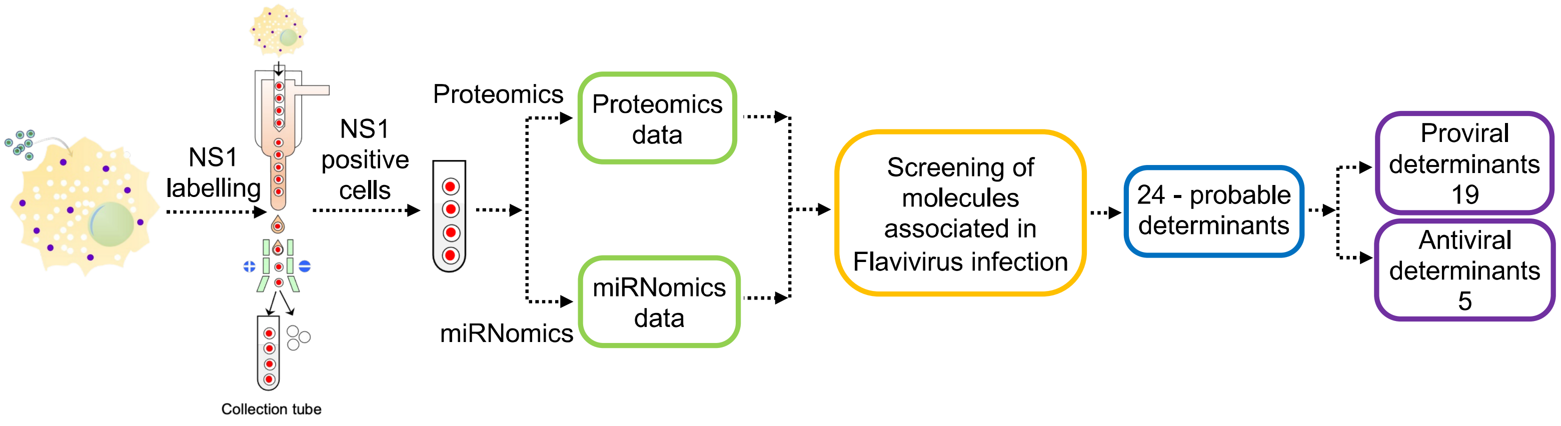

NS1
labelling
NS1
positive
cells
Collection tube
Proteomics
miRNomics
Proteomics
data
miRNomics
data
Screening of
molecules
associated in
Flavivirus infection
24 - probable
determinants
Proviral
determinants
19
Antiviral
determinants
5

A

```
Target: MK585066.1:10395-10977
length: 583
MiRNA: ssc-miR-133a-3p
length: 21

mfe: -25.7 kcal/mol
p-value: 1.000000e+00

Position: 553
target 5' G        GGU     AAGAACACA     U  3'
           UAGCUGGU   GAGG           GGAUC
           GUCGACCA   UUCC           CCUGG
miRNA  3'        AC                  UU 5'
```

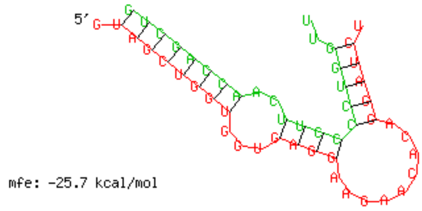


B

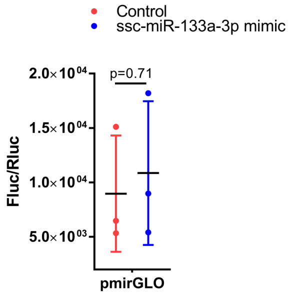


C

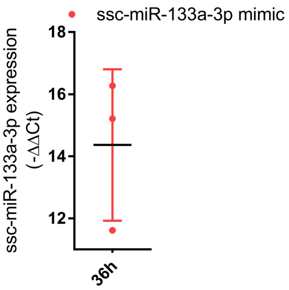


D

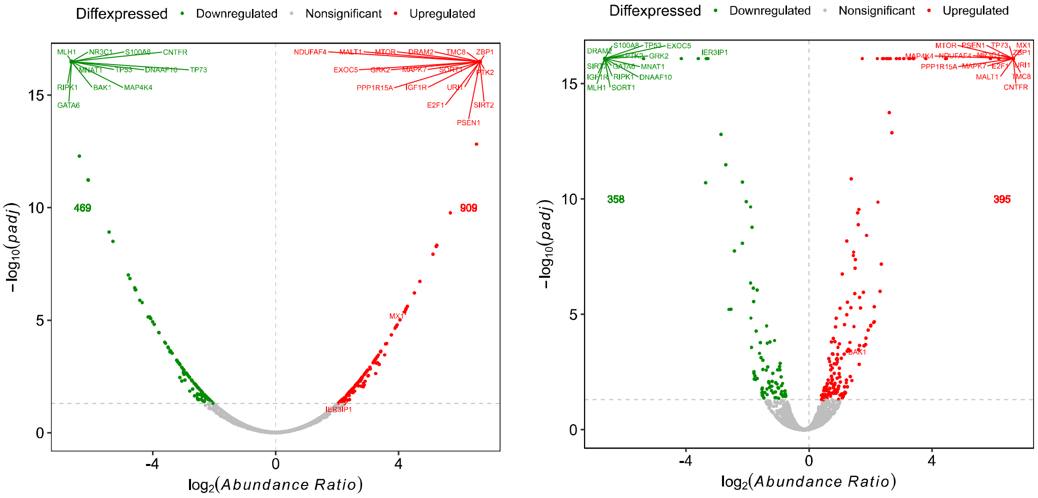


E

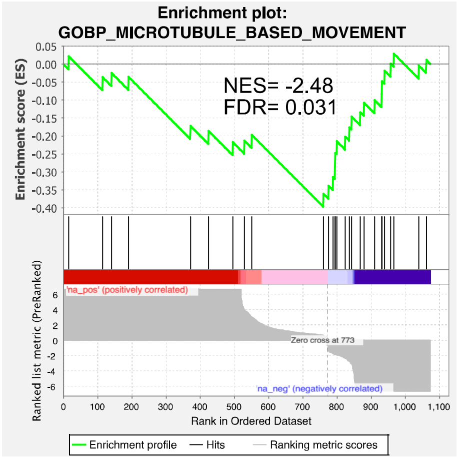

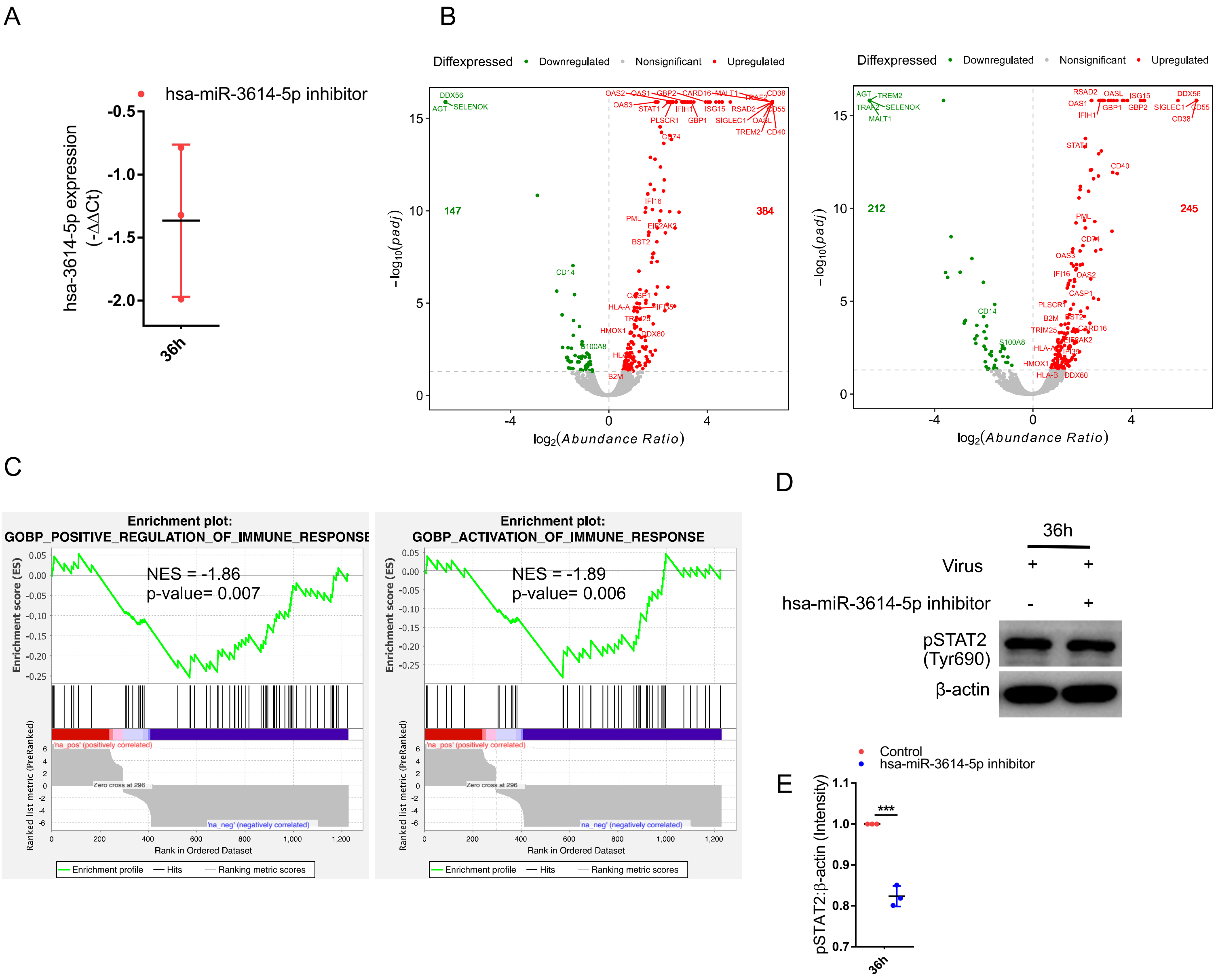
A
hsa-miR-3614-5p inhibitor
hsa-3614-5p expression (-ΔΔCt)
-0.5
-1.0
-1.5
-2.0
36h
B
Diffexpressed
Downregulated
Nonsignificant
Upregulated
147
384
212
245
−log10(padj)
log2(Abundance Ratio)
C
Enrichment plot:
GOBP_POSITIVE_REGULATION_OF_IMMUNE_RESPONSE
NES = -1.86
p-value= 0.007
Enrichment plot:
GOBP_ACTIVATION_OF_IMMUNE_RESPONSE
NES = -1.89
p-value= 0.006
Enrichment score (ES)
Ranked list metric (PreRanked)
'na_pos' (positively correlated)
Zero cross at 296
'na_neg' (negatively correlated)
Rank in Ordered Dataset
Enrichment profile
Hits
Ranking metric scores
D
36h
Virus
hsa-miR-3614-5p inhibitor
pSTAT2 (Tyr690)
β-actin
E
Control
hsa-miR-3614-5p inhibitor
pSTAT2:β-actin (Intensity)
***
36h
